# Excitation Spillover from PSII to PSI Measured in Leaves at 77K

**DOI:** 10.1101/2024.04.18.590023

**Authors:** Ichiro Terashima, Riichi Oguchi, Kimie Atsuzawa, Yasuko Kaneko, Masaru Kono

**Affiliations:** Department of Biological Sciences, School of Science, The University of Tokyo, Tokyo, 113-0033, Japan; Institute of Sustainable Agro-ecosystem Services, Graduate School of Agricultural and Life Sciences, The University of Tokyo. Nishi-Tokyo 188-0002, Japan; Comprehensive Analysis Center for Science, Saitama University, Saitama, 338-8570, Japan; Department of Natural Science, Faculty of Education, Saitama University, 338-8570, Japan

**Author notes:** Institute of Molecular Biology, College of Life Sciences, National Chung Hsing University, Taichung City 40227, Taiwan (Ichiro Terashima). Botanical Gardens, Osaka Metropolitan University, Katano 576-0004, Japan (Riichi Oguchi). Astrobiology Center, National Institutes of Natural Sciences, Mitaka 181-8588, Japan (Masaru Kono). Corresponding authors Ichiro Terashima (,), Masaru Kono; Corresponding Author Ichiro Terashima, Institute of Molecular Biology, College of Life Sciences, National Chung Hsing University, 145 Xingda Rd., South Dist., Taichung City 40227, Taiwan.

**Keywords:** fluorescence induction, grana, state transitions, sun/shade chloroplasts

## Abstract

Heterogeneous distribution of PSI and PSII in thick grana in shade chloroplasts is argued to hinder spillover of chlorophyll excitations from PSII to PSI. To examine this dogma, we measured fluorescence induction at 77K at 690 nm (PSII) and 760 nm (mainly PSI) in the leaf discs of *Spinacia oleracea*, *Cucumis sativus* and shade tolerant *Alocasia odora*, grown at high and low light, and quantified their spillover capacities. PSI fluorescence (*F*I) consists of the intrinsic PSI fluorescence (*F*I*_α_*) and fluorescence caused by excitations spilt over from PSII (*F*I*_β_*). When *F*I *and F*II parameters between State 1 and State 2, induced by weak far-red and blue light, were compared, PSII maximum fluorescence (*F*II*_m_*) and *F*I*_β_* were greater, and *F*I*_α_* was smaller in State 1 and thereby the spillover ratio, *F*I*_β_*/(*F*I*_α_ +F*I*_β_*) or *F*I*_β_*/*F*I*_m_*, was greater in State 1. Since the leftover *F*II*_m_* was found to be about 10% of total *F_m_* at 760 nm, all the data were corrected. Even after the correction, the spillover ratio in *F*I*_m_* in State 1 ranged from 21 to 32%, and the spillover ratios were comparable irrespective of growth light conditions. Although extensive grana in low light grown plants would suggest that PSII and PSI are too separated for spillover, in *A. odora* chloroplasts, the ratio of non-appressed thylakoid membranes/total thylakoid membranes was little affected by growth light and more than 40%. Abundant non-appressed thylakoids would contribute to efficient spillover.

## Introduction

Land plants are exposed to various light fluctuations. Sudden changes in the photon flux density (PFD) from low to high levels often cause photoinhibition in PSII and PSI (for a recent review, see Kono et al. 2024).

Plants have evolved several mechanisms to protect photosystems from photoinhibition. Leaf and chloroplast movements are effective (Kasahara et al. 2002, Pastenes et al. 2005), yet these are beyond the present scope. The non-photochemical quenching (NPQ), by which chlorophyll excitations are dissipated as heat, has been studied most intensively. Hereafter, chlorophyll and chlorophyll excitation are abbreviated Chl as E*. Changes in the protein structure and the energy levels of the pigments in the PSII antenna systems due to protonation of PsbS protein and conversion of violaxanthin to zeaxanthin, via antheraxanthin, are responsible for NPQ (Bassi and Dall’Osto 2021). It has been proposed that, during NPQ induction, some light harvesting Chl protein complexes (LHCIIs) are clustered and functionally separated from the PSII core complexes, and that E*s in such antenna clusters decay emitting heat (Ruban et al. 2012). It takes 5 to 10 min to induce NPQ mostly at ordinary temperatures. Full conversion of violaxanthin to zeaxanthin takes some more time (Bassi and Dall‘Osto 2021). Conversely, when the PFD level is lowered, NPQ is relaxed and the mechanism that dissipates E*s as heat is turned off. The relaxation processes take several minutes as well.

For safe and efficient photosynthesis in the fluctuating light, NPQ should be induced and relaxed rapidly. Kromdijk et al. (2016) overexpressed three components involved in the induction and relaxation processes of NPQ, violaxanthin de-epoxidase, PsbS, and zeaxanthin epoxidase, in *Nicotiana tabacum* (tobacco). When gas exchanges of such leaves were assessed in the laboratory using red and blue LEDs as actinic light, the NPQ relaxation processes were accelerated. In the field, where light fluctuates, these plants grew better than the wild type. Three lines of *Arabidopsis thaliana* overexpressing these three components showed faster NPQ induction and relaxation in the laboratory. However, neither of these lines grew better than the wild type in a natural light greenhouse. In an artificial fluctuating light in a growth chamber, these lines grew less than the wild type (Garcia-Molina and Leister 2020). Although the discrepancy of the results between *N. tabacum* and *A. thaliana* has not been resolved, the strong induction of NPQ might exert a negative effect on photosynthetic production *in A. thaliana* under their experimental conditions (Garcia-Molina and Leister 2020).

We recently found that far-red (FR) light accelerates NPQ relaxation in *A. thaliana* (Kono et al. 2020). It appears that the H^+^/K^+^ antiporter in thylakoid membranes (KEA3) is activated by FR light and accelerates NPQ relaxation (Kono et al. unpublished observation). The results of the field experiments by Kromdijk et al. (2016), therefore, would reflect this FR light effect on the rapid NPQ relaxation, because FR light is abundant in natural light.

The state transitions realise balanced distribution of E*s between PSII and PSI by adjusting allocation of mobile LHCIIs between these photosystems. The LHCII movements from PSII to PSI and from PSI to PSII are caused by the phosphorylation and dephosphorylation of LHCIIs and the PSII core, and the responsible kinases have been identified. The state transitions were once thought to play roles in protecting PSII from photoinhibition in high light (Horton and Lee 1985, Anderson and Andersson 1988, Bellafiore et al. 2005). However, it has been revealed that these kinases are inactivated in high light *in vivo* (Rintamäki et al. 1997, Pursiheimo et al. 1998). Moreover, the mutant deficient in one of these kinases, *stn7*, grew well in continuous high light. Since *stn7* did not grow well in the fluctuating light, LHCII phosphorylation may play a role in acclimation to the fluctuating light (Grieco et al. 2012).

The spillover denotes transfer of E*s from PSII to PSI. Analyses of fluorescence induction at 77K revealed that induction of PSI fluorescence occurred synchronously with that of PSII. Spillover occurs even in the absence of closed PSII. When PSII reaction centres close, more E*s are transferred to PSI. This mechanism was first described by Murata (1969). Soon after, theoretical analyses were also made (Kitajima and Butler 1975a, b, Kitajima 1976, Strasser and Butler 1977a, b). Since the spillover prevents accumulation of E*s around the closed PSII reaction centres, the intersystem crossing of the excited Chls in the singlet state (^S^Chl*s) to those in the triplet state (^T^Chl*s) around or in the reaction centres is suppressed. Although the energy level of ^T^Chl* is lower than that of ^S^Chl*, its lifetime is longer than that of ^S^Chl* by 3 orders of magnitude (Khorobrykh et al. 2020). Thus, once ^T^Chl*s are formed, the chance of ^S^O_2_ formation increases. Hence, the spillover alleviates PSII photoinhibition. E*s transferred from PSII to PSI oxidize P700 to P700^+^, the key component for protection of PSI from photoinhibition (Miyake 2020). Therefore, the spillover protects not only PSII but also PSI from photoinhibition (Terashima et al. 2021, Kono et al. 2024).

The extent of NPQ is smaller in shade plant leaves than in sun plant leaves, due to the smaller pool size per leaf area of xanthophyll cycle carotenoids in shade leaves (Demmig-Adams and Adams 2006). Shade plants in the canopy understories are often exposed to sun-flecks (Pearcy 1990) or sun patches (Smith and Berry 2013). To avoid photoinhibition due to such drastic changes in PFD, spillover would be more efficient than NPQ, because spillover occurs instantaneously while induction of NPQ requires several minutes. Shade plants, therefore, might rely more on the spillover than sun plant leaves to avoid photoinhibition.

After the classical studies for spillover quantification (Kitajima and Butler 1975a, b, Kitajima 1976, Strasser and Butler 1977a, b), there had been few attempts to quantify spillover. Recently, researchers resumed to study spillovers in various organisms using time-resolved fluorescence spectroscopy (Chukhutsina et al. 2019). Megacomplexes have been attracted attention, and the spillover is claimed to occur in the PSI-PSII megacomplexes, which include the PSI, PSII and LHC complexes (Yokono et al. 2015, Ifuku 2023, Kim et al. 2023). If this is true, transfer of E*s would occur in the single membrane because the megacomplex exists in a single thylakoid membrane.

Thylakoid membranes are categorized into appressed and non-appressed membranes. The appressed membranes occur inside the grana in contact with other appressed membranes, while the non-appressed membranes directly face the stroma, forming the inter-grana thylakoids or the grana surface membranes. In the thylakoid membranes, there are some major protein complexes, and their distributions are heterogeneous (Anderson and Andersson 1988). PSII complexes are mainly in the appressed membranes, while PSI complexes and H^+^-ATPases are mainly in the non-appressed membranes. Since the PSII complexes in the appressed membranes are located distant from the PSI complexes in the non-appressed membranes, it would be unlikely for E*s to be transferred from PSII to PSI across the membranes. Yet, the transfer of E*s across the membranes was claimed to occur based on measurements of electrogenicity in thylakoid preparations (Trissl et al. 1987).

It is known that the chloroplast morphologies are different between sun and shade plants. Grana in shade plant chloroplasts are generally thicker and contain more thylakoids (Anderson 1986, Björkman 1981). Similar trends have been reported for sun and shade leaves of the same species (Anderson 1986, Björkman 1981), and for the adaxial and abaxial parts in the same leaf (Skene 1974, Terashima et al. 1986). These morphological features imply that spillover would occur more easily in sun-type chloroplasts that have thinner grana. Chow et al. (1988) showed an electron micrograph of gigantic grana in a leaf of low-light grown *Alocasia macrorrhiza*, a shade-tolerant Araceae species. The number of thylakoids per granum exceeded 100. These impressive grana suggest that the spillover of E*s from PSII to PSI might be hampered.

In a recent paper, in which the spillover in *Pinus sylvestris* (Scots pine) leaves was detected by the time-resolved fluorescence analysis, the authors claimed that the increase in the spillover was accompanied by the decrease in the number of thylakoids per granum or by thylakoid de-stacking (Bag et al. 2020). Yokono et al. (2019) surveyed the existence of the megacomplexes in various green plants and found that sun plants accumulate more PSI-PSII megacomplexes than shade plants. They found a tendency that PSI of sun plants had deeper traps (low-energy Chls) to receive excitation energy and showed PSI fluorescence maxima at wavelengths longer than 730 nm. Unfortunately, typical shade-tolerant plants were not included in their plant materials (Table 1 of Yokono et al. 2019). Based on the spectroscopic evidence of the deep-trap Chl *a* in leaves of a shade-tolerant *Alocasia odora*, we proposed that the shade-tolerant plants also have PSI-PSII megacomplexes and, due to the deep-trap Chl *a*, spillover of the excitation energy from PSII to PSI would contribute to formation of P700^+^ in high light (Terashima et al. 2021, Kono et al. 2024).

**Table 1.**
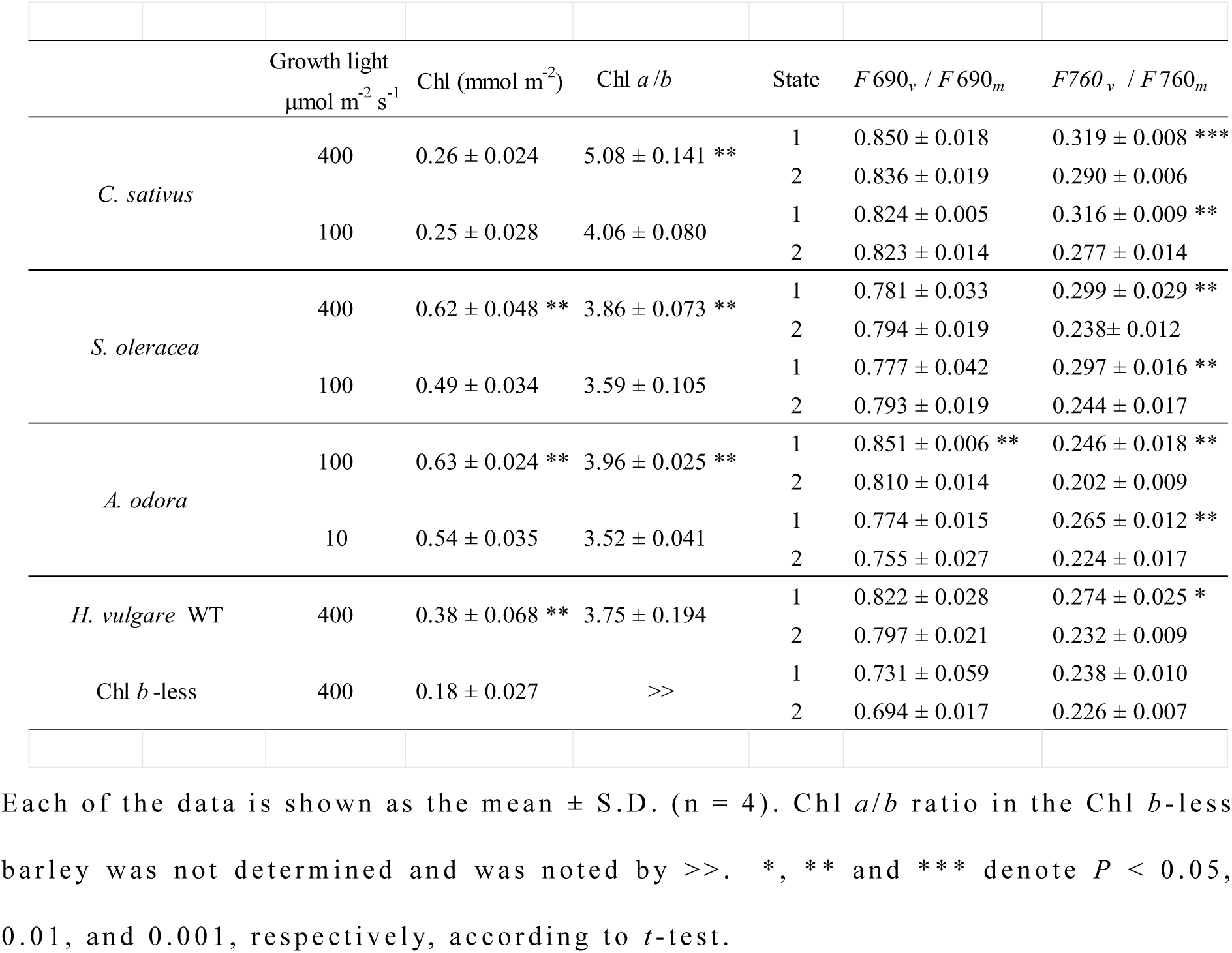
Chl content and Chl *a/b* of the samples and the effects of State transitions on *F*690*_v_*/*F*690*_m_* and *F*760*_v_*/*F*760*_m_*.

|  | Growth light<br>μmol m<sup>-2</sup> s<sup>-1</sup> | Chl (mmol m<sup>-2</sup>) | Chl *a/b* | State | *F* 690<sub>v</sub> / *F* 690<sub>m</sub> | *F*760<sub>v</sub> / *F*760<sub>m</sub> |
| --- | --- | --- | --- | --- | --- | --- |
| *C. sativus* | 400 | 0.26 ± 0.024 | 5.08 ± 0.141 ** | 1 | 0.850 ± 0.018 | 0.319 ± 0.008 *** |
|  |  |  |  | 2 | 0.836 ± 0.019 | 0.290 ± 0.006 |
|  | 100 | 0.25 ± 0.028 | 4.06 ± 0.080 | 1 | 0.824 ± 0.005 | 0.316 ± 0.009 ** |
|  |  |  |  | 2 | 0.823 ± 0.014 | 0.277 ± 0.014 |
| *S. oleracea* | 400 | 0.62 ± 0.048 ** | 3.86 ± 0.073 ** | 1 | 0.781 ± 0.033 | 0.299 ± 0.029 ** |
|  |  |  |  | 2 | 0.794 ± 0.019 | 0.238± 0.012 |
|  | 100 | 0.49 ± 0.034 | 3.59 ± 0.105 | 1 | 0.777 ± 0.042 | 0.297 ± 0.016 ** |
|  |  |  |  | 2 | 0.793 ± 0.019 | 0.244 ± 0.017 |
| *A. odora* | 100 | 0.63 ± 0.024 ** | 3.96 ± 0.025 ** | 1 | 0.851 ± 0.006 ** | 0.246 ± 0.018 ** |
|  |  |  |  | 2 | 0.810 ± 0.014 | 0.202 ± 0.009 |
|  | 10 | 0.54 ± 0.035 | 3.52 ± 0.041 | 1 | 0.774 ± 0.015 | 0.265 ± 0.012 ** |
|  |  |  |  | 2 | 0.755 ± 0.027 | 0.224 ± 0.017 |
| *H. vulgare* WT | 400 | 0.38 ± 0.068 ** | 3.75 ± 0.194 | 1 | 0.822 ± 0.028 | 0.274 ± 0.025 * |
|  |  |  |  | 2 | 0.797 ± 0.021 | 0.232 ± 0.009 |
| Chl *b*-less | 400 | 0.18 ± 0.027 | >> | 1 | 0.731 ± 0.059 | 0.238 ± 0.010 |
|  |  |  |  | 2 | 0.694 ± 0.017 | 0.226 ± 0.007 |

Each of the data is shown as the mean ± S.D. ( $n = 4$ ). Chl $a/b$ ratio in the Chl $b$ -less barley was not determined and was noted by >>. \*, \*\* and \*\*\* denote $P < 0.05$ , 0.01, and 0.001, respectively, according to $t$ -test.

In the present study, we grew sun species such as *Spinacia oleracea* (spinach) and *Cucumis sativus* (cucumber), and a shade-tolerant species, *Alocasia odora* at different light levels. *Hordeum vulgare* (barley), a wild type and a Chl *b*-less mutant lacking most LHCIIs, were also grown. We measured the induction of PSII fluorescence at 690 nm and that of mostly PSI fluorescence at 760 nm in leaf discs of these plants frozen at 77 K. We divided PSI fluorescence into the fraction corresponding to the PSI fluorescence caused by E*s spilt over from PSII and the intrinsic PSI fluorescence. The spillover ratios were compared among the species and between the growth light levels. A substantial spillover was detected in the leaves of low light grown *A. odora*. Moreover, the spillover was not necessarily greater in high light grown plants. The barley *b*-less mutant also showed considerable spillover. Effects of NPQ induction and P700^+^ formation on the intrinsic PSI fluorescence were also investigated using low light grown *A. odora*. Since the differences in ultrastructure of between the chloroplasts from high and low light grown *A. macrorrhiza* (Chow et al. 1988) were intriguing, we quantitatively analyzed the thylakoid ultrastructure of *A. odora* to elucidate the interrelationship between the ratio of non-appressed thylakoid membranes to total thylakoid membranes, thylakoid stacking, and the spillover. During the present study, we found substantial leftover PSII fluorescence at 760 nm. We tried to quantify the leftover PSII fluorescence and its effect on the estimation of the spillover. The effect of excitation of PSI fluorescence by PSII fluorescence was also considered.

## Results

### Fluorescence induction

The leaf disc on wet paper was illuminated with LEDs peaked at 720 nm (PSI light) or 470 nm (PSII light) at a photon flux density of 10 or 5 μmol m^−2^ s^−1^ for at least 30 min to allow transitions to State 1 or State 2. The disc was then placed in an aluminum cup, kept in the dark for 1.5 to 2 min, and frozen to 77 K. Induction of fluorescence excited by blue light at 460 nm at 50 μmol m^−2^ s^−1^ was measured either at 690 nm or 760 nm for PSII fluorescence (*F*690) or mostly PSI fluorescence (*F*760) at 1 ms time intervals for 30 s. *F*690 and *F*760 both showed typical inductions (Fig. 1A). It took about 5 s to attain *F_m_* levels. When *F*760(*t*) was plotted against *F*690(*t*), a linear line was obtained (Fig. 1B). See Table 1, for the plant materials. For the measurement system, see Figs. S1 and S2.

**Fig. 1.**
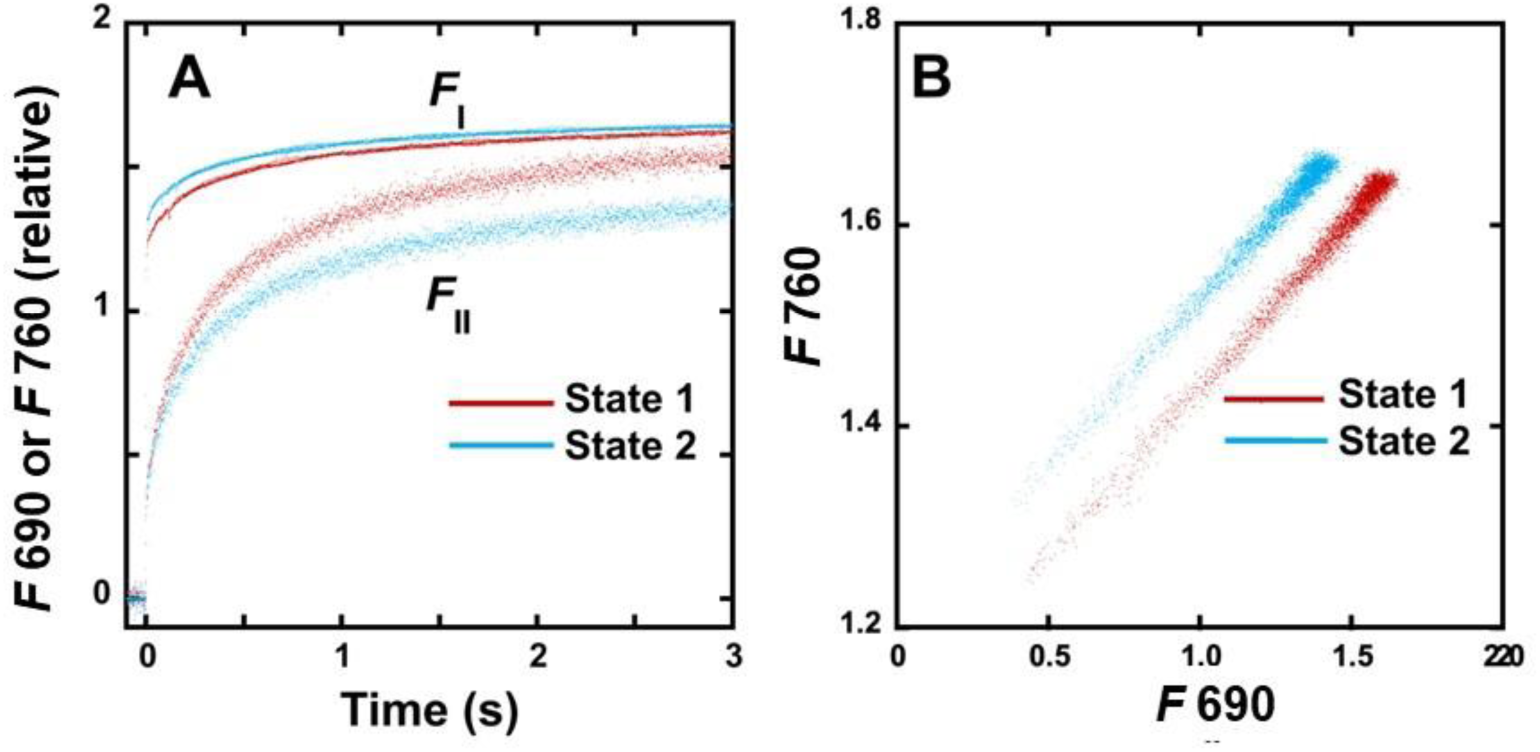
Induction curves of *F*690 and *F*760 and the relationships between *F*690 and *F*760 in LL-grown *A. odora*. A, PSII fluorescence (*F*690) and mostly PSI fluorescence (*F*760) induction measured at 690 nm and 760 nm in leaf discs at 77K of LL-grown *A. odora*. Leaf discs were illuminated with PSI light at 720 nm inducing State 1 or PSII light at 470 nm inducing State2, at the PFD of 5 μmol m^−2^ s^−1^, for at least 30 min, kept in the dark for 1.5 to 2 min and frozen in liquid N_2_. The fluorescence induction excited by the measuring/actinic light from a 460 nm LED pulse-modulated at 100 kHz was recorded. The PPFD level of the measuring/actinic light at the leaf disc was 50 μmol m^−2^ s^−1^. B, *F*760 plotted against *F*690. *S. oleracea* and *A. odora* leaf discs.

For simplicity, let us first assume that fluorescence signals detected at 690 nm and 760 nm were exclusively from PSII and PSI, and that the induction of *F*760 is attributed to the spillover of E*s from PSII (Kitajima and Butler 1985, Butler and Kitajima 1985, Kitajima 1986, Strasser and Butler 1987a, b). *F*760(*t*) is, then, expressed as:

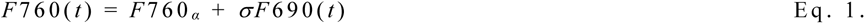

where σ is the spillover coefficient, which is equal to the ratio of variable fluorescence at 760 nm to that at 690 nm, *F*760*_v_*/*F*690*_v_*. When *F_m_* was attained:

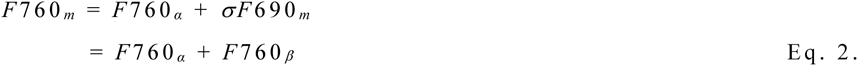

Then the proportion of the spillover in PSI fluorescence was calculated as *F*760*_β_* / *F*760*_m_* or *F*760*_β_* / (*F*760*_α_* + *F*760*_β_*). For a diagrammatical explanation of the relationships between *F*760 and *F*690, see Fig. S3.

The effects of growth light and State transitions on *F*690*_v_*/*F*690*_m_* and *F*760*_v_*/*F*760*_m_* are shown in Table 1. Except for one case of the HL grown *A. odora*, the differences in *F*690*_v_*/*F*690*_m_* depending on the States were not statistically significant, while *F*760*_v_*/*F*760*_m_* was greater in State 1. The effects on *F*690*_m_*, *F*760*_α_*, and *F*760*_m_* are shown in Table S1. In these calculations, we assumed that there was no leftover PSII fluorescence (*F*II) at 760 nm. When the data for the same leaf materials were compared, *F*690*_m_* was always greater in State 1, consistent with the view of LHCII movement from PSI to PSII in State 1 light. Inversely, PSI intrinsic fluorescence level (*F*760*_α_*), intrinsic PSI fluorescence, was greater in State 2. *F*760*_m_* is the sum of *F*760*_α_* and that is caused by the spillover from PSII, *F*760*_β_*. *F*760*_α_* was greater in State 2, while *F*7 60*_β_* would be greater in State 1, which resulted in the situation where there was no consistent effect of the State on *F*760*_m_.* These trends are relevant even when the leftover *F*II at 760 nm was considered (see Table 3, Fig. 4 and Table S2).

We also examined the induction time course (Table S1). Mobile LHCIIs would move from PSII to PSI in State 2 light and *vice versa* in State 1 light. Then, faster induction of *F*690 was expected to occur in the State 1, because antenna size per PSII reaction centre would increase in State 1. When HL and LL leaves were compared, Chl *a*/*b* was lower in LL leaves (Table 1), indicating that LHCIIs were more abundant in LL leaves. Thus, the induction would be faster in LL leaves. Both of these trends were seen (Table S1).

### Effects of the leftover PSII fluorescence (FII) at 760 nm

Hereafter, we use *F*I and *F*II to denote PSI and PSII fluorescence. Fluorescence emission spectra of the PSI particles at 77K (Lamb et al. 2018 and the references therein) indicate that there may be no PSI fluorescence (*F*I) at 690 nm. On the other hand, the PSII particles show some *F*II at 760 nm. When the PSI complexes having low energy level sinks are present together, however, E*s in the PSII complexes would be more preferentially transferred to such sinks. Thus, in the samples of high integrity, such as gently prepared thylakoids or leaf discs, E*s in PSII could be more efficiently transferred to PSI.

To assess the leftover *F*II at longer wavelengths beyond the *F*I peak, we measured fluorescence at wavelengths ranging from 690 to 780 nm. If there is *F*II at long wavelengths, *F_v_*/*F_m_* will increase with the decrease in PSI fluorescence, because *F*II*_v_*/*F*II*_m_* is greater than *F*I*_v_*/*F*I*_m_*. We also measured fluorescence emission spectra of *F_m_* in the leaf discs with a photodiode-array spectrophotometer. In these measurements, the leaves of *C. sativus*, *S. oleracea* and *A. odora* were used. The leaf discs were frozen after illumination of the State 1 light, because *F*II and thereby possible leftover *F*II would be enhanced in State 1.

Fig. 2A shows emission spectra of *F_m_* in the State 1. *C. sativus* and *A. odora* showed peaks at 740 nm, while the peak of *S. oleracea* occurred at 737 nm. The spectra for *F_v_*/*F_m_* in the State 1 are shown in Fig. 2B. At 690 nm, *F_v_*/*F_m_* in *S. oleracea* was low, indicating some chronic photoinhibition. These materials were different from those shown in Table 1 (see Materials and Methods). In all these samples, *F_v_*/*F_m_* decreased towards 750 nm and then *F_v_*/*F_m_* increased, indicating presence of leftover *F*II.

**Fig. 2.**
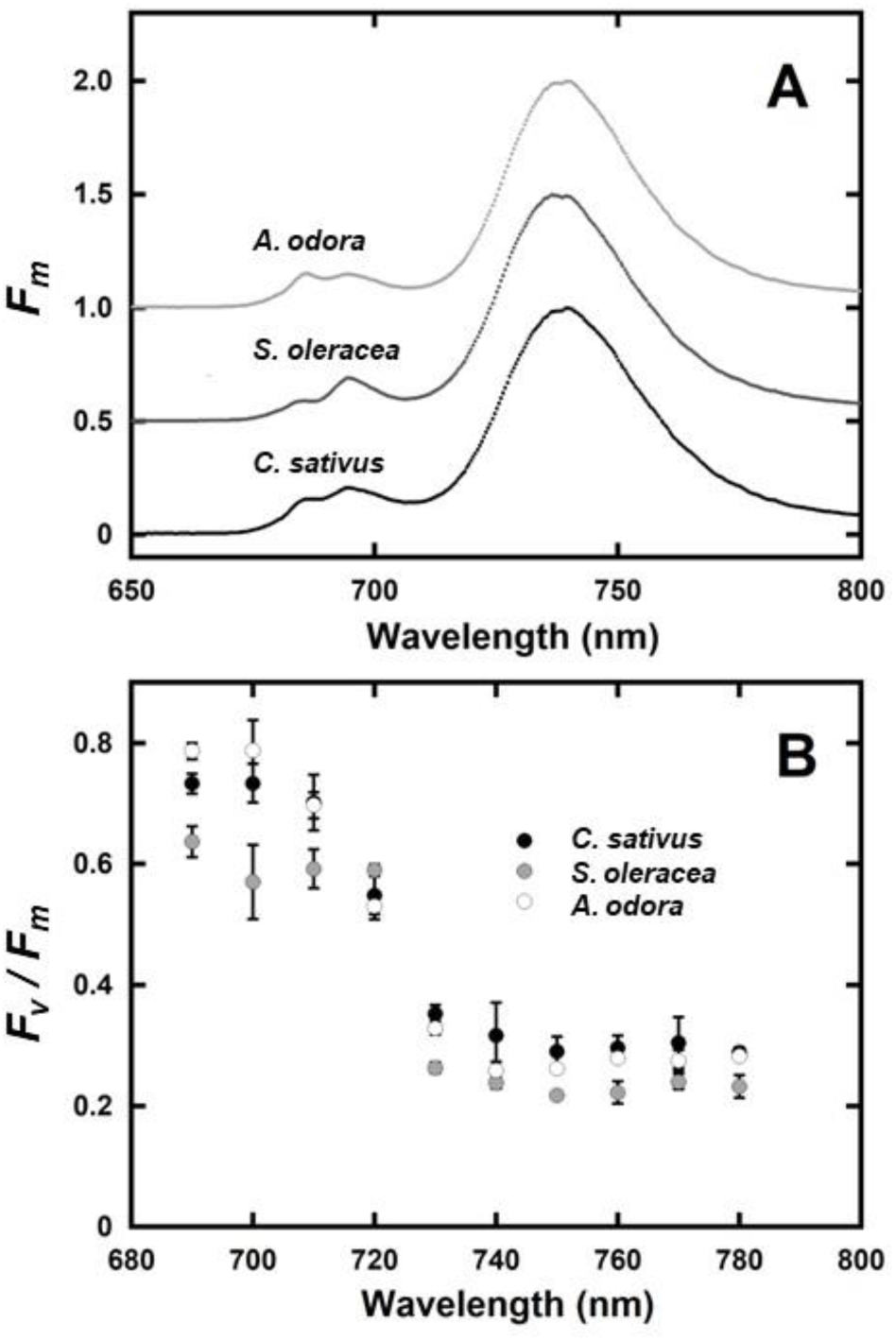
*F_m_* spectra (A) and *F_v_*/*F_m_* spectra (B) for the wavelength range from 690 to 780 nm in *C. sativus*, *S. oleracea* and *A. odora* leaf discs. A, Emission spectra of *F_m_*, excited by blue light from an LED peaked at 450 nm, were measured using a photodiode array spectrophotometer (C10083CAH, Hamamatsu Photonics, Japan). The *F_m_* spectra were not corrected for the sensitivity of the photomultiplier. B, *F_v_*/*F_m_* in the leaf discs were measured using the band pass filters (Asahi-spectra). For both A and B, the leaf discs were treated in State 1 light at 720 nm at 10 μmol m^−2^ s^−1^ at least for 30 min, kept in dim light or in the dark for 1.5 to 2 min, and frozen at 77K.

Since we used leaf discs, reabsorption of fluorescence by Chl distorted the spectral shape at short wavelengths. However, the fluorescence data at the long wavelengths can be used for quantitative analyses. Because many PSII fluorescence data of isolated LHCII and the PSII particles, except for those with obvious contamination of PSI components, show monotonous decreases towards the long wavelengths (Lamb et al. 2018 and the references therein), we assumed that the leftover *F*II*_m_* can be expressed as a linear function of *λ* for a narrow range (750 nm <λ < 780 nm). Then, *F*I*_m_*(*λ)* was expressed as *F_m_*(*λ*)-*F*II *_m_* (*λ*). *F_v_* / *F_m_* at *λ* was also expressed with *F_m_*(*λ*), *F*II*_m_*(*λ*), *F*II*_v_*/*F*II*_m_* and *F*I*_v_*/*F*I*_m_* (for details, see the supplem entary text 1). Assuming *F*II*_v_* /*F* II*_m_* was equal to *F_v_* /*F_m_* at 6 90 nm, we obtained a set of three independent variables, the slope and the intercept of the linear function expressing *F*II*_m_* (λ), and *F*I*_v_*/*F*I*_m_*, which minimiz ed the residual sum o f squares o f the differenc es (RS S) b etween the modelled and measured *F_v_* /*F_m_* values at 750, 760, 770 and 780 nm. The *F*I*_v_*/*F*I*_m_* values and the ratio s of *F*II*_m_* to total *F*_m_ at 760 nm, thus obtained, are shown in Table 2. We us ed all the measured *F_v_*/*F_m_* values for the calculations. Probably b ecaus e o f scatt ering of the data, the *F*I*_v_*/*F*I *_m_* valu e obtained for *C. sativus* was not statistically significant. When the mean *F_v_*/*F_m_* values for four wavelengths were used, however, a statistically significant value was obtained. For *A. odora* and *S. oleracea*, all the data were used. At 760 nm, *F*II*_m_* was estimated to b e 6 % of *F_m_* for *A. odora* and 1 1 % for *S. oleracea* and *C. sativus*. For the fitting o f the data, see Fig. S4. Although chang es of *F_v_*/*F_m_* with wav elength were not simple, the modelled values followed the measured values well.

**Table 2.** *F*I*_v_*/*F*I*_m_* giving the least SSR and the leftover ratio of *F*II*_m_*/*F_m_* at 760nm.

|  |  |  |  |  |  |  |  |  |
| --- | --- | --- | --- | --- | --- | --- | --- | --- |
| | $FI_v/FI_m \pm \text{sem}$ | | | | $F_v/F_m$ | | $FI_v/FI_m$ | $FII_m/F_m$ |
| | | | | | 690 nm | 760 nm | $F760_v/F760_m$ | at 760 nm |
| *C. sativus* | 0.155 | ± | 0.098 | ns | 0.733 | 0.290 | 0.534 | 0.234 |
| mean values | 0.235 | ± | 0.017 | * | 0.733 | 0.290 | 0.810 | 0.110 |
| *S. oleracea* | 0.171 | ± | 0.030 | *** | 0.637 | 0.222 | 0.770 | 0.109 |
| *A. odora* | 0.247 | ± | 0.024 | *** | 0.787 | 0.279 | 0.885 | 0.059 |

Spillover ratios are shown in Fig. 3 and Table 3. The data in Fig. 3 were calculated assuming no *F*II leftov er at 760 nm. For the spillov er ratio s as suming 0, 5, 10, and 20 % *F*II*_m_*/*F_m_* at 7 60 nm, see Table 3. As exp ected from *F*760*_v_*/*F*760*_m_* data in Table 1, spillover ratios were consistently greater in State 1 light. When HL and LL discs were compared, the levels were comparable in *S. oleracea*. But in *C. sativus* and *A. odora*, the values were significantly greater in LL discs. The leaves of shade tolerant *A. odora* showed the value of more than 0.3 in State 1. The *b*-less barley mutant also showed considerable spillover above 0.3. It is noteworthy that there were no effects of the State light on fluorescence data in the *b*-less barley, in which LHCIIs are mostly absent.

**Fig. 3.**
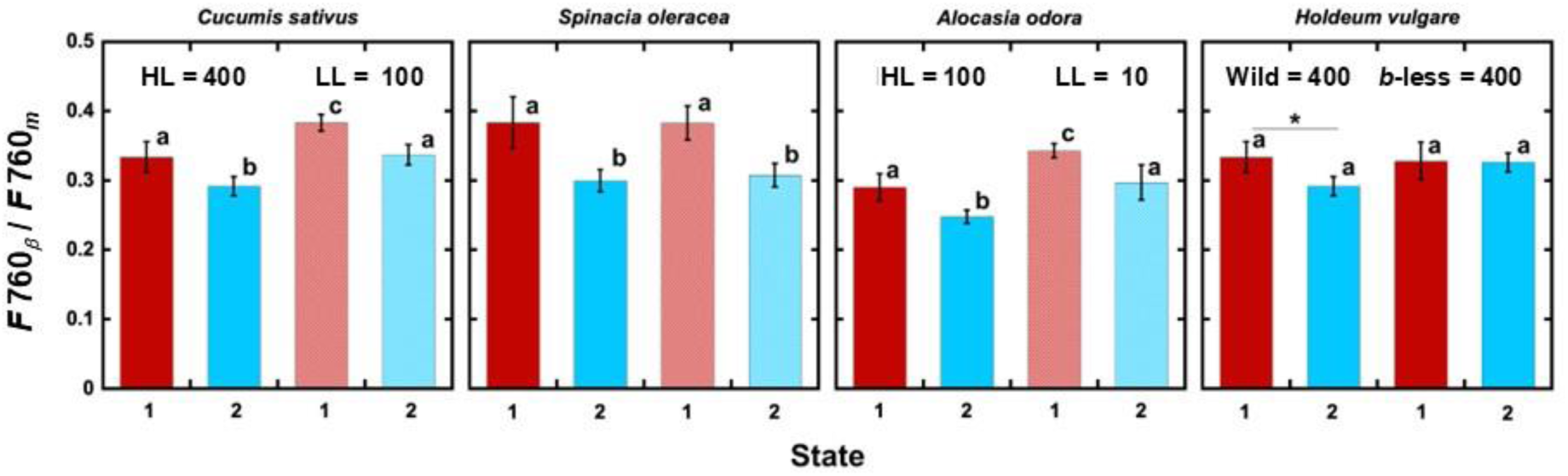
Spillover ratio (F760_β_/F760_m_) in leaf discs in the State 1 and State 2. 1 and 2 on the abscissa denote State 1 and State 2, induced by PSI and PSII light at 10 μmol m^−2^ s^−1^ at least for 30 min. For LL *A. odora*, PSI or PSII light was at 5 μmol m^−2^ s^−1^. HL and LL denote high and low growth light. 400, 100, and 10 denote growth PPFD levels in μmol m^−2^ s^−1^. Different alphabets denote statistically significant differences (*P* < 0.05) by ANOVA (Tukey-Kramer test). For HL and LL *H. vulgare*, *t*-test was used (* denotes *P* <0.05). For the data in this figure, no leftover *F*II*_m_* at 760 nm was assumed. For the effects of the *F*II leftover, see Table 3.

**Table 3.** Effects of 0, 5, 10 and 20% *F*II*_m_*/*F_m_* at 760 nm on *F*I*_v_*/*F*I*_m_* and spillover ratio (*F*I*_β_*/*F*I*_m_*)

| | Growth light<br>($\mu\text{mol m}^{-2} \text{s}^{-1}$) | State | $F690_v/F690_m$ | $FI_v/FI_m$ | | | | Spillover ratio | | | |
| --- | --- | --- | --- | --- | --- | --- | --- | --- | --- | --- | --- |
|  |  |  |  | 0% | 5% | 10% | 20% | 0% | 5% | 10% | 20% |
| *C. sativus* | 400 | 1 | 0.850 | **0.319** | 0.291 | **0.260** | 0.186 | **0.375** | 0.306 | **0.306** | 0.219 |
|  |  | 2 | 0.836 | **0.290** | 0.261 | **0.229** | 0.154 | **0.347** | 0.274 | **0.274** | 0.184 |
|  | 100 | 1 | 0.824 | **0.316** | 0.289 | **0.260** | 0.189 | **0.383** | 0.315 | **0.315** | 0.229 |
|  |  | 2 | 0.823 | **0.277** | 0.248 | **0.216** | 0.141 | **0.339** | 0.263 | **0.263** | 0.171 |
| *S. oleracea* | 400 | 1 | 0.781 | **0.299** | 0.274 | **0.245** | 0.179 | **0.379** | 0.314 | **0.314** | 0.229 |
|  |  | 2 | 0.794 | **0.238** | 0.209 | **0.176** | 0.099 | **0.287** | 0.222 | **0.222** | 0.125 |
|  | 100 | 1 | 0.777 | **0.297** | 0.272 | **0.244** | 0.177 | **0.385** | 0.314 | **0.314** | 0.228 |
|  |  | 2 | 0.793 | **0.244** | 0.215 | **0.183** | 0.107 | **0.308** | 0.231 | **0.231** | 0.135 |
| *A. odora* | 100 | 1 | 0.851 | **0.246** | 0.214 | **0.179** | 0.095 | **0.286** | 0.210 | **0.210** | 0.111 |
|  |  | 2 | 0.810 | **0.202** | 0.170 | **0.134** | 0.050 | **0.243** | 0.166 | **0.166** | 0.062 |
|  | 10 | 1 | 0.774 | **0.265** | 0.238 | **0.208** | 0.138 | **0.341** | 0.269 | **0.269** | 0.178 |
|  |  | 2 | 0.755 | **0.224** | 0.196 | **0.165** | 0.091 | **0.298** | 0.219 | **0.219** | 0.121 |
| *H. vulgare* | 400 | 1 | 0.822 | **0.274** | 0.245 | **0.213** | 0.137 | **0.331** | 0.259 | **0.259** | 0.167 |
| WT |  | 2 | 0.797 | **0.232** | 0.202 | **0.169** | 0.091 | **0.294** | 0.212 | **0.212** | 0.114 |
| *H. vulgare* | 400 | 1 | 0.731 | **0.238** | 0.212 | **0.183** | 0.115 | **0.327** | 0.251 | **0.251** | 0.157 |
| *b*-less |  | 2 | 0.694 | **0.226** | 0.201 | **0.174** | 0.109 | **0.333** | 0.251 | **0.251** | 0.157 |

When $FII_m/F_m$ at 760 nm is $\gamma$ , and if we assume $FII_v/FII_m = F_v/F_m$ at 690 nm (no $FI$ at 690 nm), then the two equations expressing $F_m$ and $F_v/F_m$ can be written:
From these, the spillover ratio can be obtained as:

Effects of *F*II leftover at 760 nm on the spillover ratios in PSI fluorescence are shown in Table 3. For a given leftover ratio, if we assum e *F*II*_v_*/*F*II_m_ is identical to *F*690*_v_*/*F*6 9 0*_m_*, *F*I*_v_*/*F*I*_m_* and the spillov er ratio (*F*I*_β_*/*F*I*_m_*) can b e calculated. When the leftover was 10%, spillover ratios in State 1 ranged from 21 to 32%. We did not examine leftovers of *F*II in either of the wild type or in the Chl *b* less mutant, the data of these materials in Fig. 3 and Table 3 should be viewed, keeping this in mind.

### Effects of NPQ

Effects of NPQ formation on spillover were examined in the leaf discs from LL *A. odora* leaf discs (Fig. 4 and Table S2). After the induction of NPQ by a white light at 700 μmol m^−2^ s^−1^ for 270 s at a room temperature of ca. 23°C, NPQ values, (*F*II*_m_* - *F*II*_m_*’)/ *F*II*_m_*’, were measured. Then, weak PSI light (at 720 nm) was given for 10 s to oxidise the acceptor side of PSII, and the leaf disc was frozen.

**Fig. 4.**
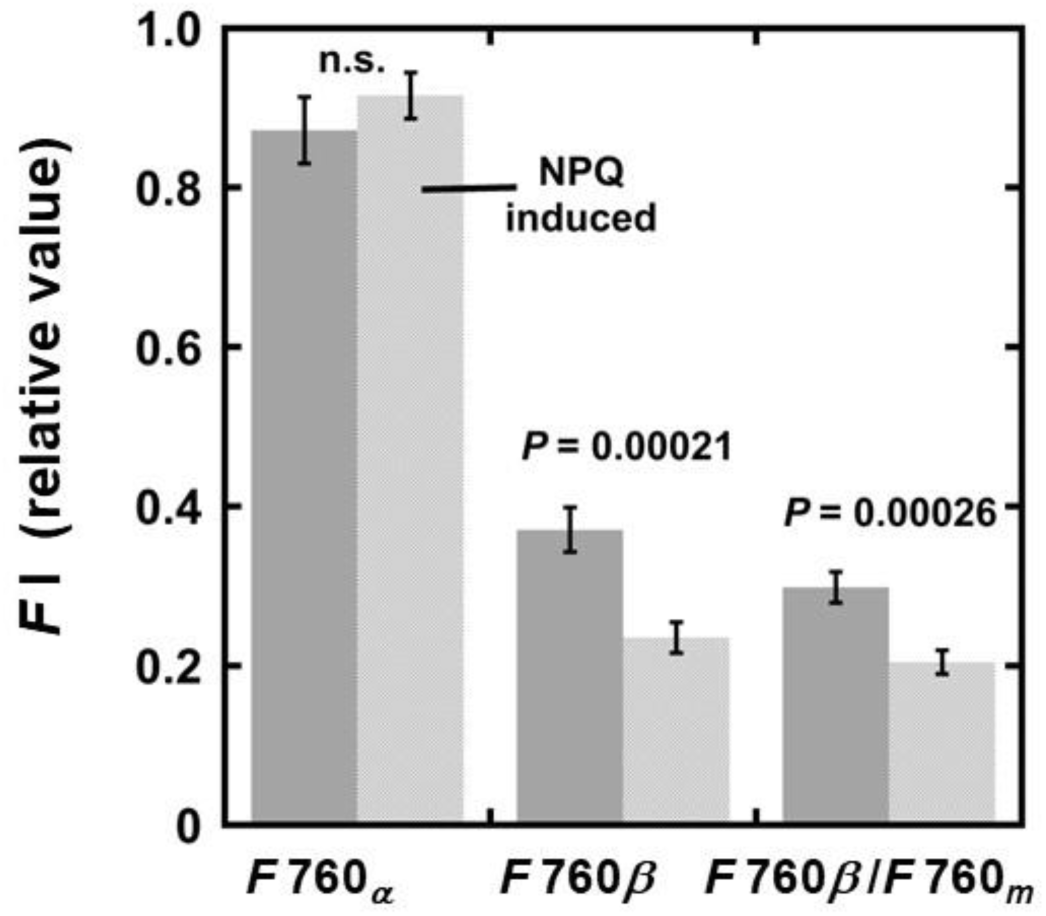
Effects of NPQ induction on spillover of excitations from PSII to PSI in leaf discs from LL-grown *A. odora*. Leaf discs from *A. odora* plants grown at 10 μmol m^−2^ s^−1^ were illuminated with a tungsten lamp at 700 μmol m^−2^ s^−1^ for 270 s and then illuminated with a 720 nm LED at 5 μmol m^−2^ s^−1^ for 10 s to oxidise the plastoquinone pool, and frozen. Inductions of *F*690 and *F*760 were measured. For the control, leaf discs treated in the dark and illuminated with the 720 nm LED at 5 μmol m^−2^ s^−1^ for 10 s were used. *P* values by *t*-test (n = 4) are shown. *F*690*_v_*/690*_m_* of the dark control and NPQ-induced samples were 0.81 ± 0.014 and 0.77 ± 0.009 (P = 0.00019, n = 4), while corresponding *F*760*_v_*/*F*760*_m_* were 0.24 ± 0.012 and 0.15 ± 0.010 (*P* = 0.00003, n=4).

Compared with sun plants, NPQ is modest in shade tolerant plants mainly due to the lower contents of xanthophyll cycle carotenoids per leaf area (Demmig-Adams and Adams III 2006). Although LL *A. odora* was used and the actinic light was moderate, NPQ levels after the NPQ induction more than 1.2 were obtained. The differences in NPQ between the samples for *F*760*_v_*/*F*760*_m_* (1.22 ± 0.119, n = 4) and for *F*690*_v_*/*F*690*_m_* measurements (1.35 ± 0.090, n = 4) were not statistically significant (*P* = 0.194, *t*-test). The data in Fig. 4, which were calculated assuming 0% *F*II*_m_* leftover at 760 nm, show smaller *F*I*_β_* and *F*I*_β_* /*F*I*_m_* in the NPQ-induced leaf discs, indicate that the spillover of E*s from PSII to PSI decreased after the NPQ induction. *F*I*_β_* /*F*I*_m_* in the dark treated control was 0.3, which resembled the value in State 2 light in LL *A. odora* (Fig. 3), as Weis (1985) pointed out for *S. oleracea*. It is also noteworthy that *F*I*_α_* was unaffected by NPQ. The effects of *F*II leftover at 760 nm on these data are shown in Table S2. The absolute value of the spillover ratio decreased with the increase in the leftover *F*II. It is noteworthy that *F*I*_α_* was calculated to be constant irrespective of the leftover ratios and NPQ induction by the equations used (see the notes for Table 3).

### Effects of P700^+^ on PSI intrinsic fluorescence

To detect PSI intrinsic fluorescence, we used a pulse-modulated measuring/actinic beam that was peaked at 700 nm and at the PFD level of 0.07 µmol m^−2^ s^−1^. Changes in the intrinsic PSI fluorescence are shown in Fig. 5. After 1 5 s from the onset of recording at 4 ms intervals, the measuring/actinic light was on. First, no noise reduction circuit was used to examine whether there was an induction in the PSI intrinsic fluorescence. Even though the measuring/actinic light was very weak, no induction was observed in both *A. odora* and *S. oleraea* (Fig. 5A and B). When an unmodulated blue light at 58 µmol m^−2^ s^−1^ was added, the fluorescence level increased to a small extent. This increase was not synchronized with the increase in P700^+^ (Fig. 5C and D). Moreover, the fluorescence levels did not respond to further increases in the PPFD level. Therefore, the increase in the ‘PSI’ fluorescence on the addition of the blue light would not be related to P700^+^. Instead, a slight increase in the ‘PSI’ fluorescence upon the blue light illumination might reflect the PSII fluorescence induction and fluorescence caused by spillover E*s from PSII to PSI. It is probable that the light used for PSI excitation slightly excited PSII. PSII reduction proceeded slowly, because the measuring/actinic beam had a narrow peak at 700 nm and was weak. Judging from the induction shown in Fig.1, PSII would be fully reduced within 5 s upon illumination of the blue light at 58 µmol m^−2^ s^−1^. Thus, the PSII fluorescence level excited by the modulated measuring/actinic light would increase. PSI fluorescence caused by the spillover from PSII contributed to this small increase. Once PSII reaction centres were all reduced, further increases in the ‘PSI’ fluorescence attributable to closure of PSII were not observed even in the stronger actinic lights.

**Fig. 5.**
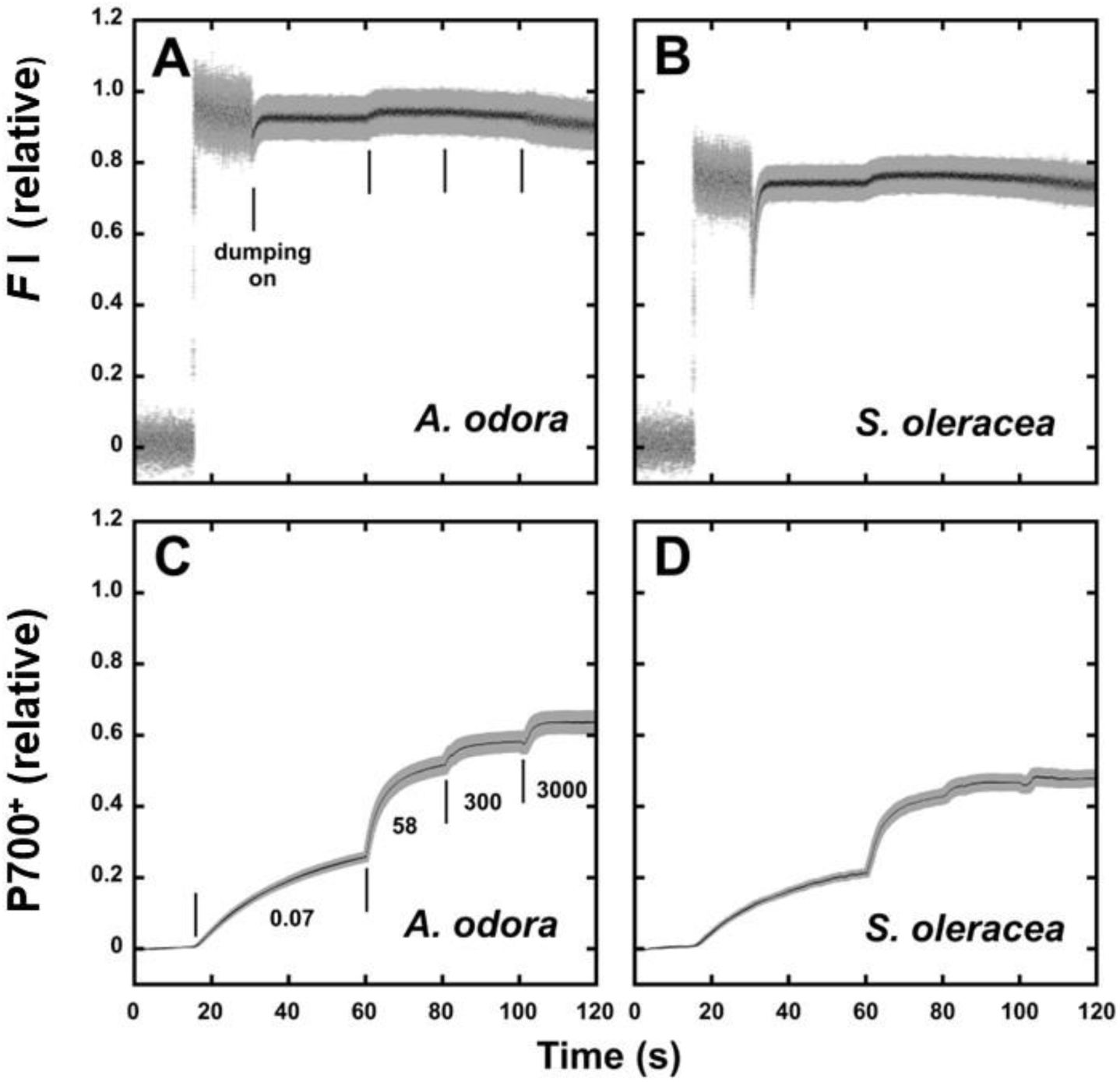
PSI intrinsic fluorescence and P700^+^ formation. A and B, Induction of PSI intrinsic fluorescence excited at 700 nm and measured in the waveband > 730 nm in the leaf discs from LL-grn *A. odora* grown at 10 μmol m^−2^ s^−1^ (A) and from spinach grown at 400 μmol m^−2^ s^−1^ (B). C and D, Formation of P700^+^ in response to 700 nm light at 0.07 μmol m^−2^ s^−1^, 470 nm blue light at 58 and 360 μmol m^−2^ s^−1^, and white incandescent light at 3000 μmol m^−2^ s^−1^.

P700^+^ increased slowly in the weak measuring/actinic light and more rapidly with 58 and 360 µmol m^−2^ s^−1^ blue light (Fig. 5C and D). A further increase in the actinic light to 3000 µmol m^−2^ s^−1^ did not cause a notable increase in P700^+^, indicating that the P700^+^ level was nearly saturated. When the ‘PSI’ traces are compared with those for the P700^+^, there were no indications of fluorescence changes with the increase in P700^+^. Upon illumination with 3000 µmol m^−2^ s^−1^ light, some decreases in ‘PSI’ fluorescence were observed both in spinach and *A. odora*.

### Effects of PSII fluorescence on PSI fluorescence

Because the *F*II emission spectrum overlaps with the absorption band of Chl *a*, some *F*II is absorbed by long waveband Chl *a* molecules in PSI. Then, *F*I would be caused not only by the spillover E*s from PSII to PSI but also directly by *F*II. In this section, the leftover of *F*II at 760 nm will not be considered for simplicity.

In our measuring system, the measuring/actinic light was illuminated from one side of the leaf disc and fluorescence emitted from the same side was measured (Fig. S1). Chls were excited with the pulse-modulated blue light at 460 nm and *F*II and *F*I were measured at 690 and 760 nm. Using a simple model, we calculated 460 nm light absorbed by a thin layer, emission of PSII fluorescence by this layer, re-abosorption of this fluorescence by the rest of the leaf, and emission of PSI fluorescence due to the reabsorption, in this order. Then, the ratio of the *F*I excited by *F*II to the *F*I directly driven by the 460nm light was calculated (see the supplementary text 2 and Fig. S5). The calculations indicate that this ratio should increase with the Chl concentration per leaf area or the Chl concentration in the thylakoid suspension (Fig. S7). If excitation of PSI by *F*II is substantial, then the ‘apparent’ *F*I*_v_*/*F*I*_m_* should increase with the Chl concentration because *F*II*_v_*/*F*II*_m_* is greater than *F*I*_v_*/*F*I*_m_*. We examined this possibility by plotting *F*760*_v_*/*F*760*_m_* of the suspensions of spinach thylakoids of varying Chl concentrations in a 3 mm thick aluminum cup (see Fig. 6A and Fig. S1). We also plotted the *F*760*_v_*/*F*760*_m_* against the Chl content in the leaf discs used for Fig. 3 (See Table 1 for Chl contents and Chl *a*/*b* ratios of the samples). We found no dependences of *F*760*_v_*/*F*760*_m_* on the chlorophyll content in the sample (Fig. 6A and B). These results indicate that the effect of PSI excitation by *F*II was small in the present measurement. This would be attributed to low PSII fluorescence yield *in vivo*, which is a few percent (Latimer et al. 1957, Lamb et al. 2018).

**Fig. 6.**
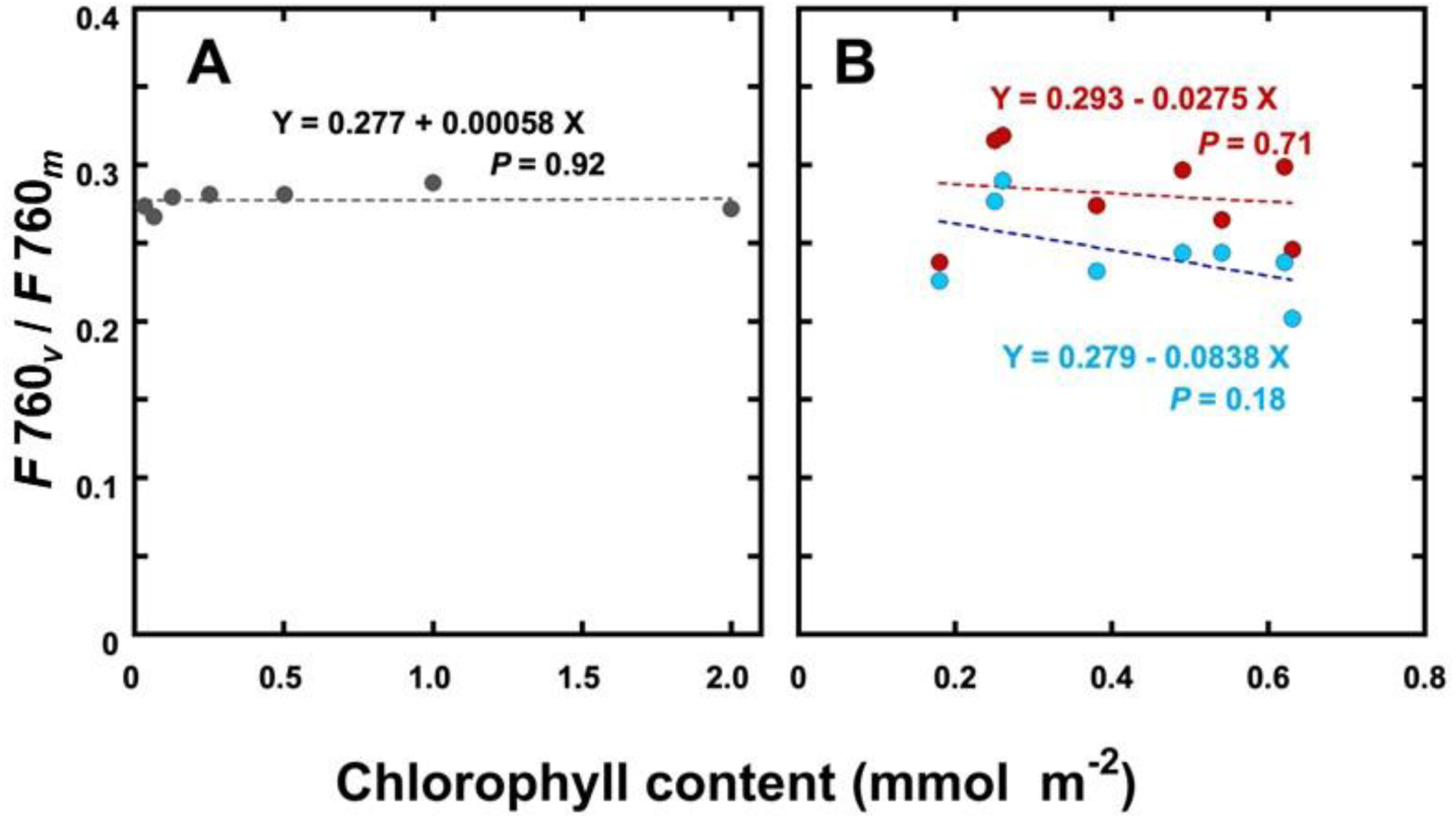
*F*760*_v_*/*F*760*_m_* plotted against the chlorophyll concentration in the spinach thylakoid suspension (A) and in the leaf discs used for Fig. 3 (B). A, Thylakoid suspensions were not pre-illuminated. The samples were kept in the dark before freezing. B, Symbols in red denote the data obtained in the State 1, while those in light blue denote the State 2. Equations are the linear regression lines. Neither of the slopes was statistically different from 0.

### Chloroplast ultrastructure of *A. odora*

In Fig. 7, electron micrographs of the chloroplasts in the leaves from the *A. odora* plants grown at 180 (HL) and 10 (LL) μmol m^−2^ s^−1^ are shown. These chloroplasts were from the first cell layers in the palisade tissues to avoid the effects of intra-leaf light gradient on thylakoid morphologies (Skene 1974, Terashima et al. 1986, Terashima and Hikosaka 1995). Thylakoid membranes per unit volume were more abundant in LL chloroplasts than in HL chloroplasts. Near the chloroplasts, peroxisomes (P) and mitochondria (M) were observed in both samples (Fig. 7 A and B). Starch grains (S) and lipid components (plastoglobules) were more conspicuous in HL chloroplasts (Fig. 7A) than in LL chloroplasts (Fig. 7B). In both samples, directions of the grana stacks were variable. Thus, both circular face views and side views of grana stacks were seen within the same sections. The diameters of grana were greater in the LL chloroplasts than in HL chloroplasts. The number of thylakoids per granum was somewhat smaller in LL chloroplasts than in HL chloroplasts. In HL chloroplasts, huge grana with numerous thylakoids were occasionally observed. In these huge grana, some thylakoids were swollen, and, in places, stackings were not tight. In both chloroplasts, there were abundant inter-grana thylakoids.

**Fig. 7.**
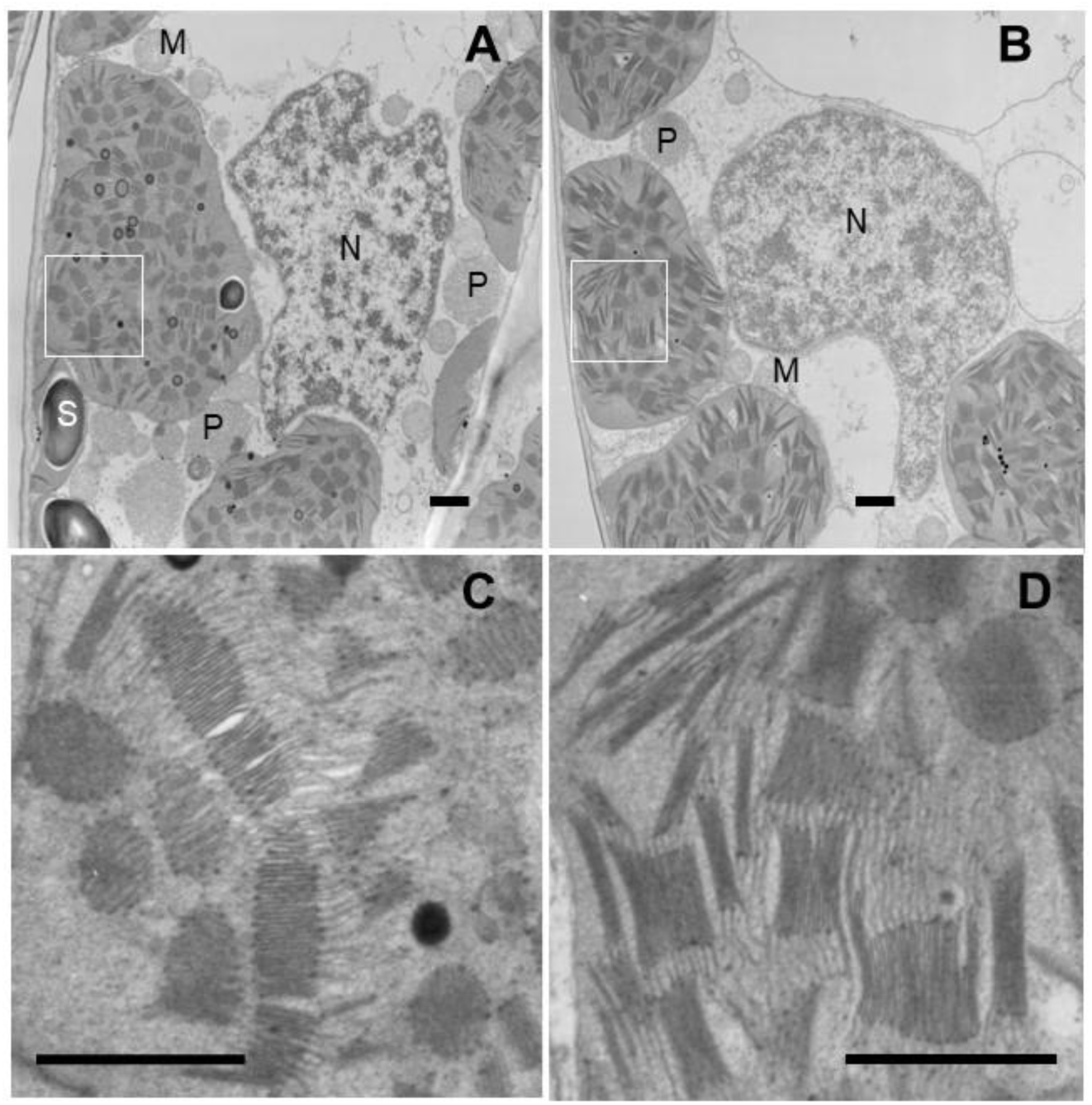
Portions of the palisade tissue cells of *A. odora* leaves grown at high light (A and C) and low light (B and D). For the electronmicroscopy study, the leaves grown at 180 and 10 μmol m^−2^ s^−1^ were used. SNear the well-developed chloroplasts, peroxisomes (P) and mitochondria (M) are observed in both samples (A and B). The square parts in A and B are enlarged in C and D. Starch grains (S) and lipid components (platoglobules) are more conspicuous in HL chloroplasts (A) than in LL chloroplasts (B). There are abundant non-appressed thylakoid membranes extending from grana stacks. Bars indicate 1 μm. The non-appressed thylakoid membranes / total thylakoid membranes in the central parts of these chloroplasts were 0.42 and 0.41 (See Figs. S9 and S10).

In Fig. 8A. the total length of thylakoid membranes per 1 μm^2^ of the ultrathin sections are shown. As already described above (Fig. 7), thylakoid membranes were more packed in the LL chloroplasts than in HL chloroplasts. The quantitative analysis of the ratios of non-appressed membrane/total thylakoid membranes (Fig. 8B) revealed that the ratios were similar between HL and LL chloroplasts: The inter-grana thylakoids were abundant in both HL and LL leaves. For the method used for the measurement of the lengths of non-appressed and appressed membranes, see Figs. S9 and S10.

**Fig. 8.**
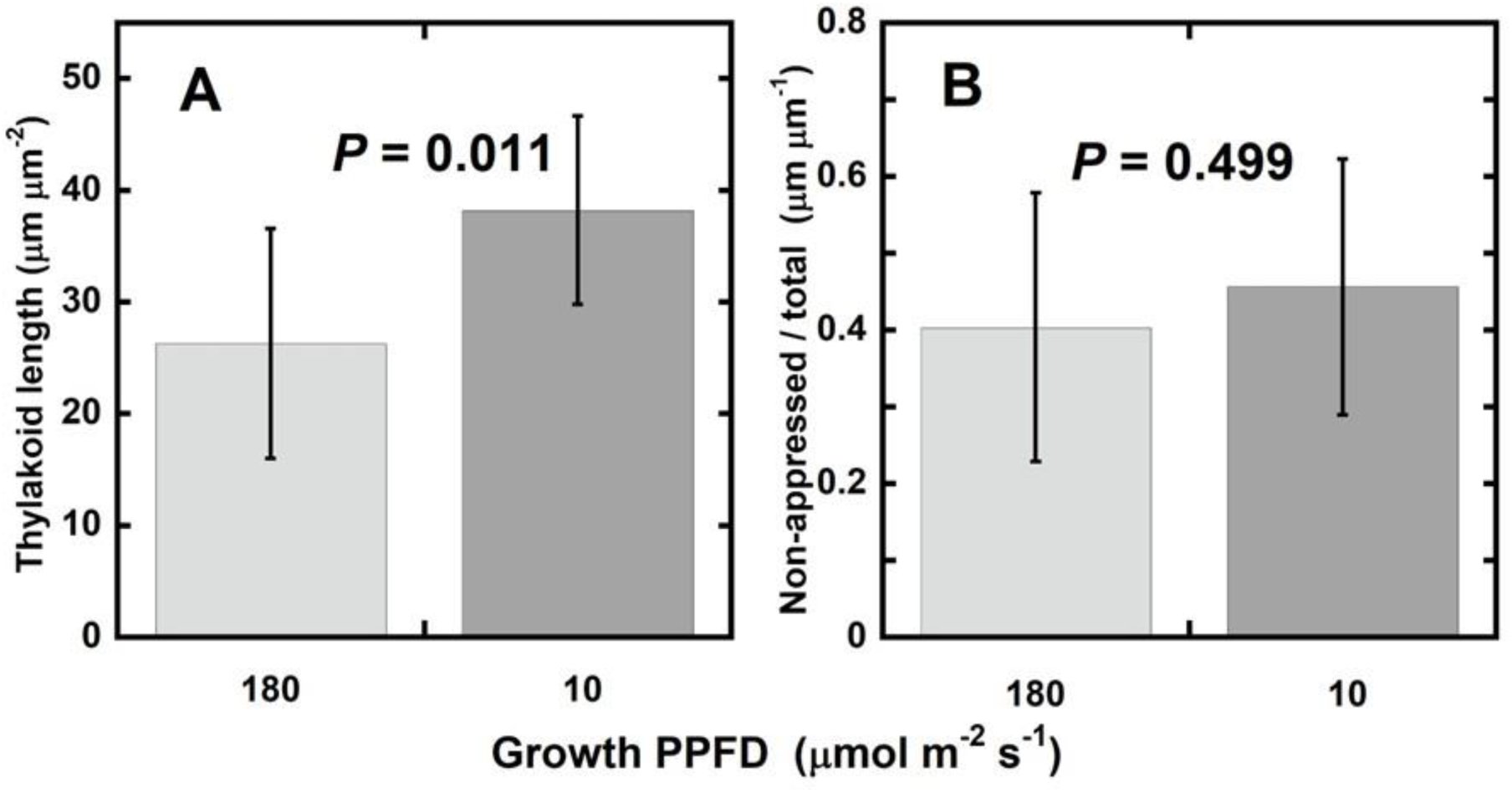
Total length of thylakoid membranes per 1 μm^2^ area (A) and the ratio of the total length of non-appressed thylakoid membranes to that of total thylakoid membranes (B). On electron micrographs of chloroplasts, 1 μm^2^ squares in which all the thylakoid membranes were in focus were chosen, and the morphometry data, total thylakoid membrane length (A) and the ratio of the total length of non-appressed thylakoid membranes to that of total thylakoid membranes (B) were obtained for leaf segments from an *A. odora* leaf grown at 180 μmol m^−2^ s^−1^ and that grown at 10 μmol m^−2^ s^−1^. The bar shows the mean for 10 squares (1 μm^2^) from 10 different chloroplasts ± SD.

## Discussion

### Quantitative analyses of the spillover

Recent studies on the spillover have been employing the time-resolved fluorescence spectral analysis (Yokono 2015, Bag et al. 2020). Although time scales of spillover and the roles of respective pigments have been clarified by this method, quantitative analyses of the quantum yield of the spillover may not be feasible. For example, it is hard to estimate how much of E*s move from PSII to PSI, although an attempt to measure the quantum yield of spillover in mixtures of cyanobacterial PSI and PSII preparations based on the steady-state and time-resolved fluorescence spectroscopy has been recently reported (Akhtar et al. 2024). Classically, slower changes of fluorescence at 77K were analyzed (Kitajima and Butler 1 975a, b, Kitajima 1976, Strasser and Butler 1977a, b). We followed these classics. We also dealt with the *F*II leftover beyond the *F*I peak, which was neglected in these classical studies.

As shown in Fig. 1, in the time scale of the present study, inductions of PSI and PSII fluorescence were synchronous as has been shown (Kitajima and Bulter 1975b). Thus, we attributed the main part of PSI induction to the spillover of excitations from PSII. We also assumed that spillover occurs in the absence of closed PSII (Kitajima and Bulter 1975 a, b, Strasser and Butler 1977b). For the diagram of these relationships between *F*I and *F*II, see Fig. S3. We assumed that *F*I*_α_* is constant, which would be supported by the data shown in Figs. 4 and 5. The mathematical equations used for the calculation of *F*I*_α_* give constant *F*I*_α_* at any leftover ratios. The constant FI*_α_* values shown in Figs. 4 and 5, thus, inversely support the relevance of our simple equations shown in the note for Table 3.

Although these classical studies neglected the F*II* leftover at the long wavelengths, we found the substantial leftover of *F*II at 760 nm. Unfortunately, we could not devise a direct method to quantify the *F_v_*/*F_m_* leftovers. However, we tried to quantify the leftovers from *F_m_* and *F_v_*/*F_m_* spectra, based on the simple assumptions (see the supplementary text 1). Our estimations of the PSII leftovers were 6% for *A. odora*, and 11% for *S. oleracea* and *C. sativus*. Probably these values should not be regarded as values specific to the species. Instead, these may be regarded as examples.

The contribution of spillover to *F*I is summarized in Fig. 3. The lowest value was recorded for the HL grown *A. odora* treated in State 2 light. However, this was still more than 0.24. The highest values approached 0.38 in State 1 light. After the correction for the 10% *F*II*_m_*/*F_m_* at 760 nm, these values were 0.17 and 15 0.32, respectively (Table 3).

One of the main aims of the present study was to examine whether the spillover occurs in LL acclimated chloroplasts. Considerable spillover occurred in the shade tolerant plant *A. odora*, and the spillover ratio in this plant was greater in the LL-leaf than in the HL-leaf. In *C. sativus* as well, the LL leaf showed higher spillover ratios, while *S. oleracea* the spillover values were similar.

When NPQ was developed, the spillover from PSII to PSI became smaller. It is also noteworthy that *F*I*_α_* was unaffected, indicating that NPQ did not develop in the PSI antennae or its effect was negligibly small in PSI. In *Chlamydomonas reinhardii*, the protonated LHCSR3 is involved in energy dissipation in both PSI and PSII antennae (Tokutsu et al. 2013, Girolomonia et al. 2019). However, its close homologs are not found in land plants.

Using the time-resolved fluorescence spectral analysis, it has been shown that the excited state of deep-trapped Chl *a* is quenched by P700^+^ in the PSI-PSII megacomplex prepared from *Arabidopsis thaliana* (Yokono et al. 2019). It is inferred that PSI fluorescence is quenched in the presence of P700^+^. For the Förster resonance transition of the excitation from the excited pigment A to B to occur, the fluorescence emission spectrum of A should overlap with the absorption spectrum of B. The probability is also proportional to the inverse of the sixth power of the distance between these pigments (Blankenship 2021). Because P700^+^ has a broad absorption band (Inoue et al. 1973), which well overlaps with the PSI fluorescence emission spectrum from red Chls, Förster resonance transition would be possible from E*s to P700^+^. When this transition occurs efficiently, quenching of PSI fluorescence with the increase in P700^+^ may be expected. However, appreciable quenching was not detected in LL-grown *A. odora* or *S. oleracea* (Fig. 5), probably because the quenching was subtle if any, due the longer distance from the red Chls to P700 (more than 6 nm) than the distance (~ 4.2 nm) in cyanobacteria (Karapetyan et al. 1999, Schlodder et al. 2005).

On the other hand, in the P700-enriched particles (Ikegami 1976) and the PSI preparation (Wientjes and Croce 2012), P700^+^ increases the PSI fluorescence level. Although Ikegami (1976) found a considerable increase in PSI fluorescence in his PSI particle, having the Chl/P700 ratio of 7, he argued that this much increase would be hidden in the thylakoids. Wientjes and Groce (2012) detected a 4% increase in PSI fluorescence with the formation of P700^+^ in their PSI preparation in which the chlorophyll/P700 ratio would be around 130. In the data shown in Fig. 5, we were not able to detect the effects of P700 oxidation level on PSI intrinsic fluorescence level. Thus, at least for the first approximation, we would be able to regard *F*I*_α_* as a constant.

PSI would be excited by PSII fluorescence. The ratio of the PSI fluorescence excited by PSII fluorescence (*FI*_←*F*II_) to the PSI fluorescence directly excited by the 460 nm light (*FI_dírect_*) increases with the Chl content per unit area (See the supplementary text 2 and Fig. S7). Then, *F*I*_v_*/*F*I*_m_* should also increase with the increase in the Chl content per area. However, this trend was not observed (Fig. 6). *FI_←Fll_/FI_dírect_* may be expressed by Eq. S14 (see the supplementary text 2), which is the product of the yield of PSII fluorescence and the ratio of the integrals. The former would be a few percent (Latimer et al. 1956, Lamb et al. 2018). The latter would be less than 1~2. Thus, *F_I←Fll_/F_I dtrect_* would be negligible.

Concerning the procedures of the present study, it had been better to measure the ‘mostly’ PSI fluorescence at 750 nm instead of 760 nm. Then, the overestimations of spillover ratios were smaller. The *F_v_*/*F_m_* spectra had to be measured first, and then the wavelength for the mostly PSI fluorescence measurement had to be determined. The system used in the present study is not ideal for such measurements. The time-resolved fluorometer that can measure the time-dependent spectral transient from *F*_0_ to *F_m_* such as used in Franck et al. (2002) or Santabarbara et al. (2019) should be used. Then, the full spectra can be measured with one leaf disc. Based on the present study, we propose the important procedures needed for accurate quantification of the spillover.

### Spillover in fluorescence quenching analyses

Fluorescence is a useful probe to analyse photosynthetic reactions (Papageorgiou and Govindjee 2004). However, in many fluorescence models, spillover is not considered. Let us assume that the initial allocation of E*s to PSII is *ψ*_ii_.

Maximum quantum yield of PSII photochemistry on absorbed PFD (*Φ*_ii_), with no energy dependent quenching (NPQ), may be expressed as follows (Kono et al. 2024). In the present consideration, we deal with *F*II_0_ and *F*II*_m_* only.

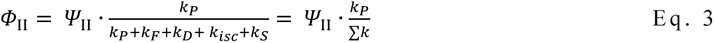

where *k’s* are the first-order rate constants: *k_P_*, photochemistry; *k_F_*, radiative (fluorescence) deexcitation; *k_D_*, thermal deexcitation to the ground state via pathways other than NPQ (non-photochemical quenching); *k_isc_*, intersystem crossing leading to formation of ^T^Chl*; and *k_S_* spillover to PSI. Let us assume that the quantum yield of *F*II*_v_*/*F*II*_m_* with fully oxidised PSI in low light is 0.8:

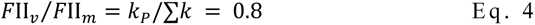

Thus,

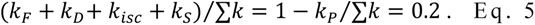

In weak light, PSI quantum yield on absorbed PFD basis, *Φ*i, is expressed as follows:

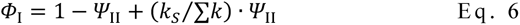

We may assume that *Φ*_II_ and *Φ*_I_ are identical in weak light for the smooth linear electron transport to occur. Then,

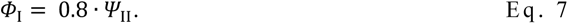

As the spillover ratios in State 1 with closed PSII, after the correction for 10% *F*II*_m_*/*F_m_* at 760 nm, ranged from 0.210 ~ 0.3 15 (Table 3):

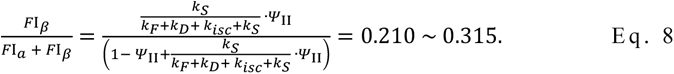

Noting Σk = 5 • *(k_F_ + k_D_ + k_lsc_* + *k_s_),* we can solve Eq. 7 and Eq. 8. Then, *ψ_II_* ranges from 0.568 to 0.577 and *k_s_/Σk* ranges from 0.040 to 0.067. In other words, even when PSII reaction centres are all open, 4 to 7% E*s initially allocated to PSII are redirected to PSI. When all the PSII reaction centres are closed, 20 to 27% of E*s initially allocated to PSII are transferred to PSI. Although the relative contribution of spillover decreases with the increase in NPQ, the present analyses confirmed that the spillover ratio is large, and it should not be neglected. It is worth pointing out that considering the lower quantum yield of PSII and spillover, the initial allocation of E*s to PSII should be much greater than 0.5, which is usually assumed in many models.

### Chloroplast ultrastructure

Figs. 7 shows chloroplast structure of *A. odora* leaves. As expected, thylakoid membrane length per unit volume was greater in the chloroplasts in the LL leaf than those in the HL leaf. By dividing these lengths by 2, the thylakoid lengths are obtained. Although some swelling was observed in HL thylakoids, assuming a thylakoid thickness of 13 nm (Kirchhoff 2019), the thylakoid volume per unit chloroplast volume, excluding inter-thylakoid spaces, was calculated as 17% or 25% in HL or LL chloroplasts.

For the Förster-type transfer to occur, the set of pigments should be located close enough. For Chls, the distance which allows 50% probability of energy transfer is calculated to be 5~10 nm (Van Grondelle et al. 1994), then the jump across the inter-thylakoid partition (~ 4 nm, according to Kirchihoff 2019) would be possible. Then, energy transfer may not be confined to within the same membrane. However, based on analyses of the decay time of fluorescence, it has been argued that the main pathway of the spillover is within the same membrane (Farooq et al. 2016). It is noteworthy that inter-grana thylakoids were extensive and that the fractions of the non-appressed thylakoid membranes were similar and more than 40% in both HL and LL chloroplasts of *A. odora*. Although the grana morphology has been attracted attention (Anderson 1986, Terashima and Hikosaka 1995) and rearrangement of grana in response to light has been shown (Anderson and Andersson 1988, Rozak et al. 2002, Kirchhoff 2019), detailed dynamics of the inter-grana thylakoids have not been examined.

The appressed thylakoid membranes may also accommodate megacomplexes in their margins. For example, the size of a PSI-PSII megacomplex of *Oryza sativa* (rice) is approximately 25 nm in its major axis (Kim et al. 2023) and the diameter of the granum ranges 400 to 600 nm (Fig. 6, see also Kirchhoff 2019). Assuming that the grana diameter is R(nm) and the megacomplexes are aligned in the margin of the thylakoid with their PSI portions outwards, the area of the thylakoid accommodating the megacomplexes is calculated to be *[R^2^* — *(R* — *25)^2^]/R_2_.* Thus, the marginal area corresponding to 23 to 16% of the appressed thylakoid membrane may accommodate the megacomplexes. Since the area of the appressed thylakoids were 60 and 54% of the total areas for HL and LL chloroplasts (Fig. 8), the membrane area that may accommodate the PSI-PSII megacomplexes would be 53% (0.40+0.60 x 0.23) and 55% (0.46 + 0.54 x 0.16) of the total membrane areas in HL and LL chloroplasts.

### Conclusions and a brief ecological consideration

The present study showed that spillover occurs in all plant materials including LL grown shade tolerant *A. odora*. The organized spillover would occur within the PSI-PSII megacomplexes in the non-appressed thylakoid membranes and in 19 the marginal areas of appressed-thylakoids. It is noteworthy that these parts would comprise more than 50% of the total thylakoid membranes in both HL and LL grown *A. odora*. In nature, shade tolerant species are usually in the green shade. In such green shade, far-red light is enriched (Kono et al. 2020, 2024). Thus, the thylakoids of these plants are in the State 1, and operate photosynthesis at the highest efficiency without NPQ. Upon sudden exposure of such leaves to high light due to sun patches (Smith and Berry 2013) or sun-flecks (Pearcy and Way 2012), E*s initially allocated to PSII would be spilt over to PSI most efficiently. Since the decreased electron flow and increased excitation flow to PSI contribute to formation of P700^+^, the safe quencher, the spillover would protect both PSII and PSI from photoinhibition by the spillover.

## Materials and Methods

### Plant Materials

*Cucumis sativus* L. (cucumber) “Nanshin,” *Spinacia oleracea* L. (spinach) “Torai,” *Alocasia odora* (G. Lodd.) Spach, and *Hordeum vulgare* L. (barley) “Gunilla, Svalöf AB N83-4001,” a wild type, and *H. vulgare* “Dornaria, chlorina-f2^2800^,” a chlorophyll *b*-less mutant, were grown in vermiculite in pots in growth chambers at 25°C. Light period was 12 h. Light was provided by a bank of white fluorescent lamps. Photosynthetic photon flux density (PPFD) levels were adjusted with the number of fluorescent tubes and/or the black shade cloth. *C. sativus*, *S. oleracea*, and *A. odora* were grown at two photosynthetic PPFD (400– 700 nm) levels. For cucumber and spinach, the PPFD levels were 400 and 100 μmol m^−2^ s^−1^, while for *A. odora*, 100 and 10 μmol m^−2^ s^−1^ were used. *H. vulgare* plants were grown at 400 μmol m^−2^ s^−1^. For electron microscopy, *A. odora* grown at 180 and 10 μmol m^−2^ s^−1^ were used. Plants were given the 1/1000 strength of the Hyponex 6–10–5 solution (Hyponex, Japan) diluted with tap water every two or three days. The relative humidity of the chamber was kept above 50%. The growth light PPFD level of 400 μmol m^−2^ s^−1^ for sun plants or crops would be sufficiently high. At 1 0 μmol m^−2^ s^−1^, where *A. odora* showed slow growth, corresponds to PPFD levels on the deep forest floor.

For the assessment of *F*II leftover at 760 nm, we grew *A. odora* and *C. sativus* outdoors in the soil in the pots. The *A. odora* plant used was the clone of the plants used in other measurements. For *C. sativus,* the same variety (‘Nanshin’) was used. We bought *S. oleracea* from a local market. These measurements were conducted in July 2024, the warmest season.

## 77K fluorescence from leaf discs

Fluorescence induction at 77K was measured with a PAM 101 fluorometer (Walz, Effeltrich, Germany). The original red LED (650 nm) used for the pulse-modulated measuring light was replaced with a blue LED (OSUB51 1 1AST, OptoSupply, Hong Kong) peaked at 460 nm. In the PAM 101 fluorometer, a photodiode was used to detect fluorescence. The original far-red filter was replaced with a 690 nm or 760 nm bandpass filter (HMX0690 or HMX0760, Asahi-Spectra, Tokyo, Japan) for PSII or mostly PSI fluorescence measurements. Leaf discs (10 mm in diameter) were cut with a leaf punch. 16 leaf discs were used for one series of measurements. For one set of measurements, four leaf discs were used, and fluorescence inductions were measured at 690 and 760 nm, respectively, in States 1 and 2. These four discs were taken from a narrow uniform part of a leaf. For *C. sativus*, *S. oleracea*, and *A. odora* 16 leaf discs were taken from one leaf, whereas, for *H. vulgare*, four sets were taken from four separate leaves. The leaf discs were placed on wet filter paper with the adaxial sides upward and illuminated with State 2 light at 470 nm from LEDs (L470, Ushio, Tokyo) or State 1 light at 720 nm from LEDs (L720, Ushio, Tokyo) at a photon flux density of 10 µmol m_-2_ s^−1^ at a room temperature of ca. 25°C. For *A. odora* leaves grown at 10 μmol m^−2^ s^−1^, the PFD level was lowered to 5 µmol m^−2^ s^−1^. After illumination for at least 30 min with either of the State 1 or State 2 LEDs, the adaxial side of leaf disc was attached to a quartz rod with silicone grease and covered with an aluminum cap and dipped in liquid nitrogen in a Dewer jar. From the cessation of the State light treatment until the freezing, all the procedures were made within 1.5 to 2 min in dim light. After the enclosure in the cup, the sample was further darkened. When the temperature of the sample in the aluminum cap was equilibrated with that of liquid nitrogen, which was detectable by a change of the sound of bubbling, recording was started. After 2 s, the measuring/actinic blue light at 50 μmol m^−2^ s^−1^ (pulse modulated at 100 kHz) was turned on. The fluorescence induction at 690 nm or 760 nm was recorded at 1 ms time intervals for 28 s on a USB data acquisition system (USB-1608FS, Measurement Computing, Norton, MA, USA). For the measurement system see Fig. S1. For transmittance spectra of 690 and 760 nm bandpass filters, see Fig. S2.

The *F_v_*/*F_m_* spectra in the wavelength range from 690 to 780 nm were obtained as described above. Emission spectra of *F_m_*, excited by blue light from an LED peaked at 450 nm, were measured using a photodiode array spectrophotometer (C10083CAH, Hamamatsu Photonics, Japan) and an optical system similar to that used for the *F_v_*/*F_m_* measurement. For details, see Terashima et al. (2021) and Fig. S1. *F_v_*/*F_m_* in the leaf discs were measured as described above using the band pass filters (Asahi-spectra). The leaf discs were treated in State 1 light at 720 nm at 10 μmol m^−2^ s^−1^ at least for 30 min.

### Effects of NPQ on the spillover

The effects of NPQ on the spillover were examined using an *A. odora* leaf grown at 10 μmol m^−2^ s^−1^ light. The leaf disc attached to a quartz rod was covered with an aluminum cap, and NPQ was induced by illuminating the leaf disc with an actinic white light from a tungsten lamp (KL1 500 lcd, Schott, Mainz, Germany) at 700 μmol m^−2^ s^−1^ for 270 s at a room temperature of ca. 23°C. Saturating flashes at 6,600 μmol m^−2^ s^−1^ for 1 s were given at 20 s intervals. The actinic light was turned off, and 10 s after, the measuring beam was turned off. At 5 s after turning off the actinic light, far-red light peaked at 720 nm at 10 μmol m^−2^ s^−1^ was illuminated for 10 s to oxidise PSII. Then, the leaf disc in the cap was frozen at 77K. When the temperature reached 77K, PSI or PSII fluorescence was measured as above.

### Effects of P700+ formation on PSI fluorescence

PSI fluorescence was obtained by preferential illumination of PSI pigments. The actinic/measuring light was a 700 nm LED (5 mm bullet type, Ushio, Tokyo, Japan) passing through a 700 nm band pass filter with 10 nm half-band width (HMX700, Asahi-spectra, Tokyo, Japan) and a 710 nm short pass filter (SVX 710). For detection of PSI fluorescence, the photodiode of the PAM 101 covered with a long pass filter (LVX 730, allowing transmission of far-red light > 730 nm) was used. Since it was necessary to use the measuring/actinic light of very narrow waveband, the PFD level at the leaf disc surface was 0.07 μmol m^−2^ s^−1^.

The leaf disc kept in the dark at least for 30 min was attached to the quartz rod, enclosed by an aluminum cup, chilled at 77 K, and its PSI fluorescence was measured. The accumulation of P700^+^ was measured with a dual-wavelength (820/870 nm) unit (ED-P700DW) attached to a PAM fluorometer (Walz) in the reflectance mode. When the two PAM101 systems were operated at the same time, the signals sometimes became noisy. Thus, we used similar leaf discs taken from the same leaf and measured the PSI fluorescence and redox state of P700 separately, but according to the same illumination time schedule.

After the onset of recording at 4 ms intervals, the measuring/actinic light for PSI at 0.07 µmol m^−2^ s^−1^ was on at 15 s. The signal damping was started to reduce the noise level at 30 s, an unmodulated blue light at 58 μmol m^−2^ s^−1^ from 460 nm LED was added at 60 s, the blue light level was increased to 360 μmol m^−2^ s^−1^ at 80 s, and, the blue light was turned off and a white light from a tungsten lamp at 3000 μmol m^−2^ s^−1^ was turned on at 100 s. When the redox state of P700 was measured, the pulse-modulated measuring light from a PAM 101 far-red LED emitting light with wavelengths ranging from 800 to 900 nm was used.

### Effects of excitation of PSI fluorescence by re-absorbed PSII fluorescence

To examine the effects of Chl concentration on PSI fluorescence excited by reabsorbed PSII fluorescence, thylakoids were prepared from the laboratory grown spinach at 400 μmol m^−2^ s^−1^ PPFD (see above) as described in Terashima et al. (2021). PSI fluorescence induction in the thylakoid suspension in a buffer containing 0.3 M sorbitol, 10 mM NaCl, 5 mM MgCl_2_ and 50 mM and Hepes-KOH (pH at 8.0) was measured at 77K in a 3 mm thick aluminum cuvette (see Fig. S1). All these procedures were made in dim light. The pulse modulated measuring/excitation light at 460 nm used was at the PPFD of 12 μmol m^−2^ s^−1^. The weaker light was used to obtain more data points for the induction phase with the diluted thylakoid suspensions. For theoretical consideration, see the supplementary text 2.

### Data processing

*F*_0_ *and F_m_ determination for leaf discs*: The recording system was turned on at 0 s, and the measuring/actinic light was turned on at 2 s. The zero (0) and the maximum fluorescence (*F*_m_) levels were obtained by averaging 2000 data points between 0 and 2 s and between 28 s and 30 s. A linear regression equation was obtained using the 50 data points, from 2.005 to 2.054 s. *F*_0_ was obtained by substituting 2 s to this equation.

*F*_m_ and *F*_0_ thus obtained were used to calculate *F*_v_ (= *F*_m_ – *F*_0_) and *F*_v_ /*F*_m_. As the increases in PSI and PSII fluorescence intensities were synchronous (Fig. 1), the variable fluorescence in PSI was assumed to be attributed to the spillover from PSII variable fluorescence.

The spillover from PSII to PSI was assumed to occur at the same efficiency regardless of whether PSII reaction centres were open or closed (Strasser and Butler (1977a, b). Based on these assumptions, and assuming that PSII and PSI fluorescence could be separately measured at 690 and 760 nm, PSI fluorescence at time *t*, *FI(t)* was expressed as:

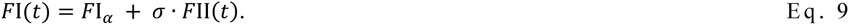

Where σ is the spillover coefficient. From this, intrinsic PSI fluorescence (*FI_a_*) was obtained:

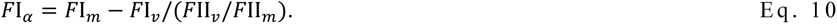

#### Induction time determination

Assuming that the PSII fluorescence induction occurred in two phases, the induction curve was fitted by the equation:

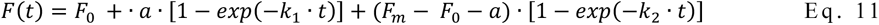

We used *F*_0_ and *F_m_,* which were obtained as above. *A*, *k*_1_ and *k*_2_ were obtained by the fitting with KaleidaGraph (Version 5.0, Synergy Software). Then, *t*_1/2_ was obtained as *t* giving *F(t) = F_0_ + (F_m_-F_0_)/2*.

#### F_0_ determination for thylakoid suspension

Since a weaker actinic/measuring light was used for thylakoid suspensions, the linear regression of 50 data points was not reliable due to the lower S/N ratio. When the number of data points was simply increased, linearity could not be assumed. Thus, we fitted Eq. 8 to the fluorescence transient for 2 s to obtain *F*_0_, *a*, *k*_1_ and *k*_2_. Because time constants *k*_2_ was always one order of magnitude smaller than *k*_1_, we further assumed that *k_2_* = 0.1 • *k_1_.* The determination coefficients obtained with this assumption was always greater than that for the equation in which *k*_1_ and *k*_2_ were independently obtained.

### Electron microscopy

The *A. odora* plants used were grown at the PPFD 180 or 10 µmol m^−2^ s^−1^. After harvesting, the leaves were kept in the dark at a room temperature for at least 30 min before fixation. Leaf segments, ca. 1 mm x 4 mm were excised and fixed immediately in 2% glutaraldehyde in 0.05 M potassium phosphate buffer (pH 6.8) and kept at the room temperature for 2 h and then at 4°C overnight. Six rinses in the buffer solution preceded postfixation with 2% OsO_4_ in the buffer at room temperature for 2 h. After dehydration in a graded acetone series, leaf materials were embedded in Spurr’s resin. The ultrathin sections (silver-gold) were poststained with uranyl acetate and lead citrate and examined with a Hitachi H-7500 transmission electron microscope at an accelerating voltage of 80 kV.

Electron-micrographs of the first cell layers of the palisade tissues were taken to avoid the effects of intra-leaf light gradient on thylakoid morphologies (Skene 1974, Terashima et al. 1986, Terashima and Hikosaka 1995). In the leaves grown at 180 or 10 μmol m^−2^ s^−1^ light, 36 or 30 chloroplasts were randomly chosen, and in these chloroplasts, 178 or 157 squares of 1 μm^2^, in which all thylakoid membranes were in focus, were selected randomly. We randomly selected 10 of these squares from 10 chloroplasts for each growth light level and thylakoid lengths were measured using Fiji/ImageJ (NIH).

The total length of the thylakoid membranes per 1 μm^2^ was also calculated as an index of thylakoids to stroma volume ratio. The abundance of non-appressed thylakoids was expressed as the ratio of the length of the non-appressed thylakoid membranes to the total thylakoid membranes. For examples of tracing the electronmicrographs, see Figs. S9 and S10.

## Acknowledgement

The barley seeds were a kind gift of Professor W.S. Chow. We thank Professor W. S. Chow, Dr. D. Y. Fan, Dr. E. Kim, Dr. M. Kitano and Dr. M. Kitajima for constructive comments. We also thank Dr. G. Shansker for drawing our attention to PSI fluorescence excited by PSII fluorescence. We are grateful to Mr. H. Hashimoto and Mr. A. Okamoto, Namoto Co., Ltd., for assisting modification of the PAM 101. We examined the leftover PSII fluorescence at 760 nm in response to the comments by the two reviewers. We thank them for this fundamental comment.

## Author Contributions

All authors contributed to this study and are responsible for the content of this paper.

## Supplementary materials

**Table S1.** *F*II induction time, *F*690_m_, *F*760_m_, and *F*760_α_.

| | Growth light<br>$\mu\text{mol m}^{-2} \text{s}^{-1}$ | State | $t_{1/2}$<br>(s) | $F690_m$ | $F760_m$ | $F760_\alpha$ |
| --- | --- | --- | --- | --- | --- | --- |
| *C. sativus* | 400 | 1 | $0.336 \pm 0.017$ | $1.58 \pm 0.075^{**}$ | $0.81 \pm 0.074$ | $0.51 \pm 0.046$ |
| | | 2 | $0.344 \pm 0.030$ | $1.41 \pm 0.027$ | $0.94 \pm 0.078^*$ | $0.62 \pm 0.054^+$ |
| | 100 | 1 | $0.255 \pm 0.017$ | $1.73 \pm 0.036^{**}$ | $1.15 \pm 0.053$ | $0.71 \pm 0.028$ |
| | | 2 | $0.279 \pm 0.016^+$ | $1.54 \pm 0.079$ | $1.18 \pm 0.053$ | $0.78 \pm 0.050^*$ |
| *S. oleracea* | 400 | 1 | $0.319 \pm 0.055$ | $1.49 \pm 0.146^+$ | $0.95 \pm 0.135$ | $0.59 \pm 0.098$ |
| | | 2 | $0.338 \pm 0.020$ | $1.30 \pm 0.082$ | $1.08 \pm 0.098$ | $0.76 \pm 0.063^*$ |
| | 100 | 1 | $0.251 \pm 0.023$ | $2.15 \pm 0.150^{**}$ | $1.35 \pm 0.103$ | $0.83 \pm 0.091$ |
| | | 2 | $0.307 \pm 0.040^+$ | $1.75 \pm 0.076$ | $1.30 \pm 0.052$ | $0.90 \pm 0.055$ |
| *A. odora* | 100 | 1 | $0.353 \pm 0.013$ | $1.40 \pm 0.030^+$ | $1.40 \pm 0.077$ | $1.00 \pm 0.077$ |
| | | 2 | $0.377 \pm 0.046$ | $1.09 \pm 0.056$ | $1.36 \pm 0.034$ | $1.03 \pm 0.030$ |
| | 10 | 1 | $0.286 \pm 0.026$ | $1.68 \pm 0.084^{**}$ | $1.67 \pm 0.071$ | $1.10 \pm 0.050$ |
| | | 2 | $0.326 \pm 0.026^+$ | $1.49 \pm 0.082$ | $1.69 \pm 0.057$ | $1.19 \pm 0.045^*$ |
| *H. vulgare* WT | 400 | 1 | $0.338 \pm 0.029$ | $1.54 \pm 0.119^{**}$ | $1.63 \pm 0.058$ | $1.09 \pm 0.059$ |
| | | 2 | $0.493 \pm 0.030^{***}$ | $1.30 \pm 0.050$ | $1.87 \pm 0.075^{**}$ | $1.32 \pm 0.063^{**}$ |
| Chl *b*-less mutant | 400 | 1 | $1.259 \pm 0.080$ | $0.41 \pm 0.070$ | $0.49 \pm 0.092$ | $0.33 \pm 0.054$ |
| | | 2 | $1.348 \pm 0.093$ | $0.44 \pm 0.062$ | $0.51 \pm 0.107$ | $0.34 \pm 0.079$ |

+, \*, \*\*, and \*\*\* denote statistically significant differences between the data ( $P < 0.1$ , $P < 0.05$ , $P < 0.01$ and $P < 0.001$ according to $t$ -test) in State 1 and those in State 2.
The effects of growth light on $t_{1/2}$ are not compared statistically, because leaf materials per se were different. However, $t_{1/2}$ values were consistently smaller in low light grown materials.

We confined ourselves to compare the absolute fluorescence values in the samples from the same leaves, because optical properties were very different between HL and LL leaves even for the same species. Accumulation response of chloroplasts would occur in State 2 light (blue light at 10 or 5 μmol m^−2^ s^−1^). This would increase *F*II*_m_*. But, in fact, *F*II*_m_* decreased in State 2. Thus, the effect of chloroplast accumulation responses was weaker than that of the State transitions.

**Table S2.** Effects of the *F*II leftover at 760 nm on the fluorescence parameters shown in Fig. 4.

|  |  | 0% | 5% | 10% | 20% |
| --- | --- | --- | --- | --- | --- |
| $FI_v/FI_m$ | DARK | **$0.241 \pm 0.0120$** | 0.211 | **0.178** | 0.099 |
|  | NPQ | **$0.152 \pm 0.0100$** | 0.119 | **0.083** | - |
| $FI_m$ | DARK | **$1.242 \pm 0.0488$** | 1.180 | **1.118** | 0.994 |
|  | NPQ | **$1.151 \pm 0.0330$** | 1.093 | **1.036** | - |
| $FI_\alpha$ | DARK | **$0.872 \pm 0.0361$** | 0.872 | **0.872** | 0.872 |
|  | NPQ | **$0.915 \pm 0.0249$** | 0.915 | **0.915** | - |
| $FI_\beta/FI_m$ | DARK | **$0.298 \pm 0.0196$** | 0.261 | **0.220** | 0.122 |
|  | NPQ | **$0.204 \pm 0.0149$** | 0.155 | **0.160** | - |

**Fig. S1.**
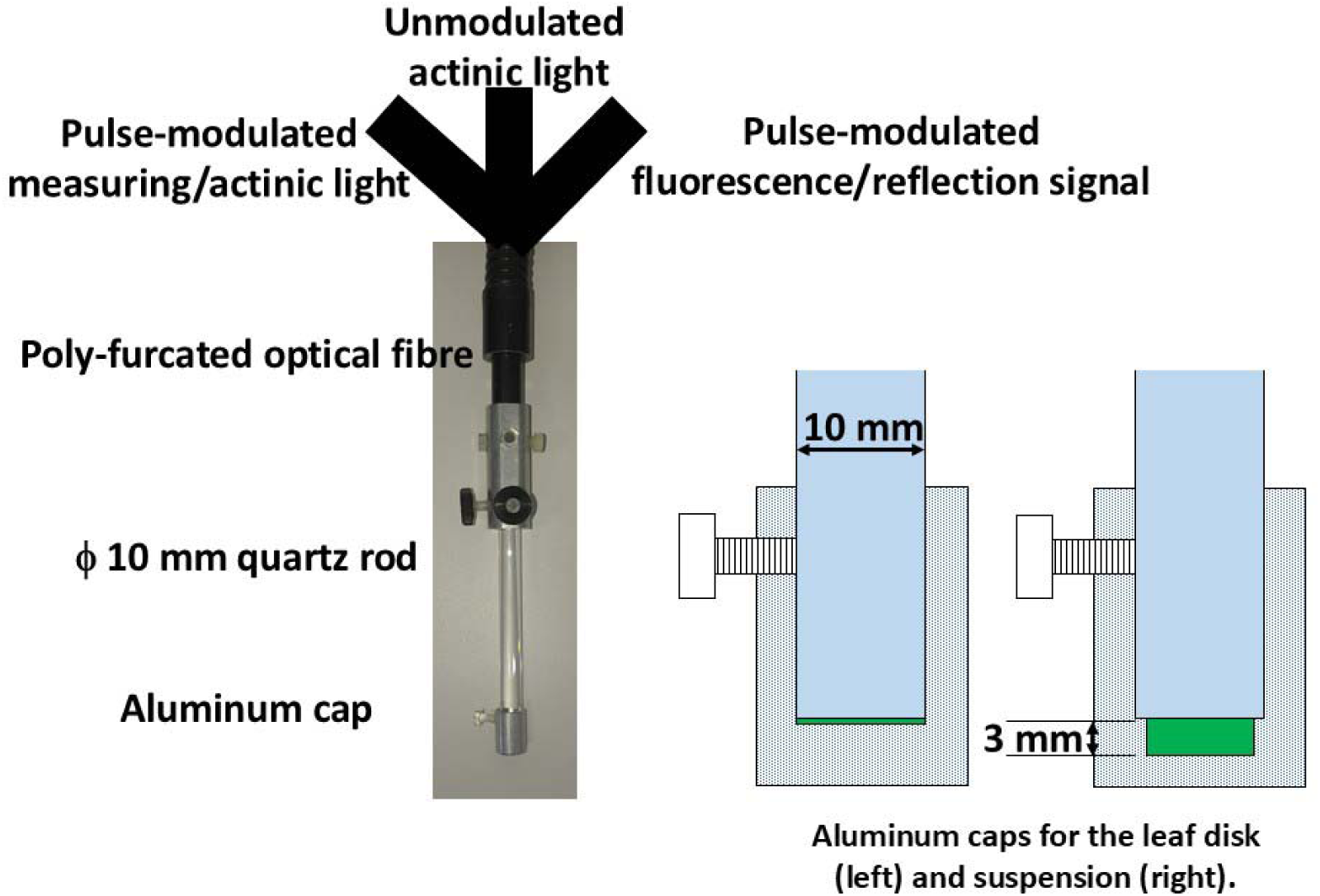
The system for *F_v_*/*F* measurement.

**Fig. S2.**
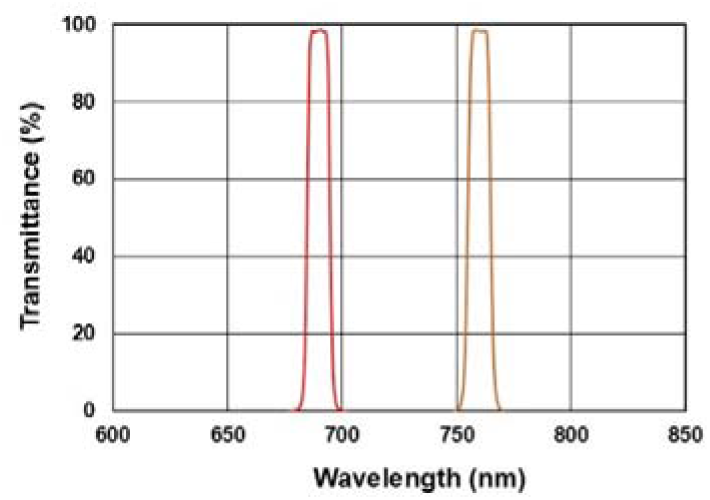
Transmittance spectra of the 690 and 760 nm bandpass filters. The half-band width was 10 nm. The tailings were small.

**Fig. S3.**
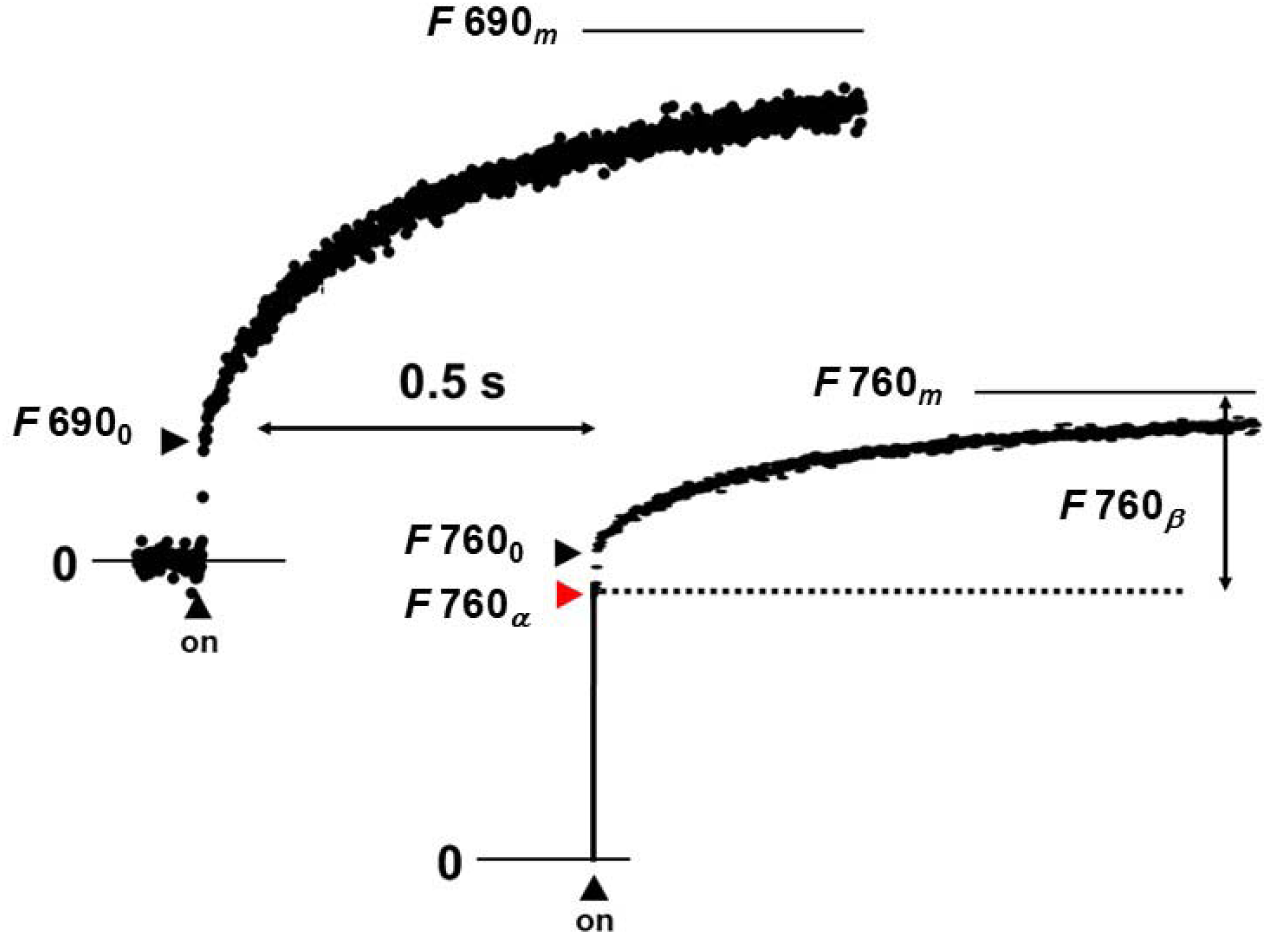
Interrelationships between *F*690 and *F*760.

When the leftover PSII fluorescence at 760 nm is absent, induction of F760 is exclusively due to the spillover from PSII. However, if there is a leftover *F*II*_m_* at 760 nm, we cannot distinguish the spillover from the contamination of *F*II.

When the leftover of *F*II*_m_*/*F_m_* at 760 nm was *γ*, and if we assume *F*II*_v_*/*F*II*_m_* = *F_v_*/*F_m_* at 690 nm (no *F*I at 690 nm), two equations can be formulated:

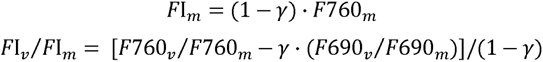

From these, the *F*I*_α_* and the spillover ratio can be obtained as:

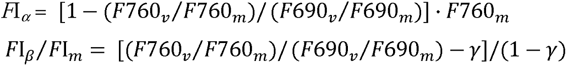

Note that *F*I*_α_* is constant at any γ

### Estimation of the leftover FII at 760 nm

For this series of measurements, we grew *A. odora* and *C. sativus* outdoors in the soil in the pots. The *A. odora* plant used was the clone of the plants used in other measurements. For *C. sativus,* the same variety (‘Nanshin’) was used. We purchased *S. oleracea* from a local market. The measurements were conducted in July 2024, the warmest season.

Emission spectra of *Fm*, excited by blue light from an LED peaked at 450 nm, were measured using a photodiode array spectrophotometer (C10083CAH, Hamamatsu Photonics, Japan) and an optical system similar to that used for the *F_v_*/*F_m_* measurement as described in Terashima *et al*. (2021). *F_v_*/*F_m_* in the leaf discs were measured as described in detail in the Materials and Methods using the band pass filters. The leaf discs were treated in State 1 light at 720 nm at 10 µmol m^−2^ s^−1^ at least for 30 min. These spectra are shown in Fig. 2 in the main text.

We assumed that the *F*II*_m_* would be expressed by a linear function for the wavelength range between 750 and 780 nm:

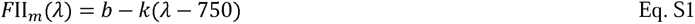

where *λ* is wavelength in nm (750 < 7 < 780). Then, the *Fm* level fluorescence at wavelength *λ* can be expressed as:

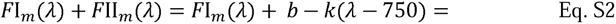

We assumed that *F*II*v*/*F*II*m* was identical to that at 690 nm, *F*690*v*/*F*690*m*. Because *F*I*v*/*F*I*m* is unknown, *Fλ_v_/Fλ_m_* is expressed as:

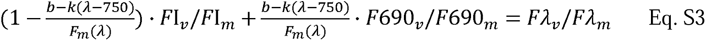

We obtained the set of *b*, *k*, *FI_v_/FI_m_* that minimized the residual sum of squares (*RSS)* of the differences between the predicted and measured *F_v_/F_m_* values at four wavelengths:

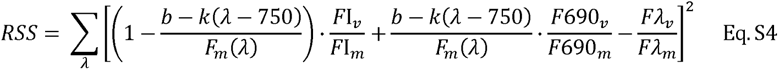

The *F*_m_ spectra shown in Fig. 2 in the main text were not corrected for the sensitivity of the photomultiplier. However, we corrected the values using the data provided by the manufacturer (Hamamatsu photonics). Calculations were made with R statistical software (version 4.2.2; R Foundation for Statistical Computing; available from http://www.R-project.org). The function *nls* was used to obtain *b, k* and *FI_v_/FI_m_* by fitting nonlinear relationship among wavelength, *Fm* and *Fv*/*Fm* with the least-squares method. We measured three *F_v_*/*F_m_* values for each wavelength for each species. The three independent variables were estimated using all these values. For *S. oleracea* and *A. odora*, *FI_v_/FI_m_* significantly significant values were obtained, whereas for *C. sativus*, the value was not statistically supported probably due to scattering of the *FI_v_/FI_m_* values. When the mean *Fv*/*Fm* value for each wavelength was used, a statistically significant *FI_v_/FI_m_* was obtained (Table 3 in the main text). Fitting of these models to the measured values are shown in Fig. S4. When statistically significant values were used, the changes in *F_v_/F_m_* with *λ* were well traced.

**Fig. S4.**
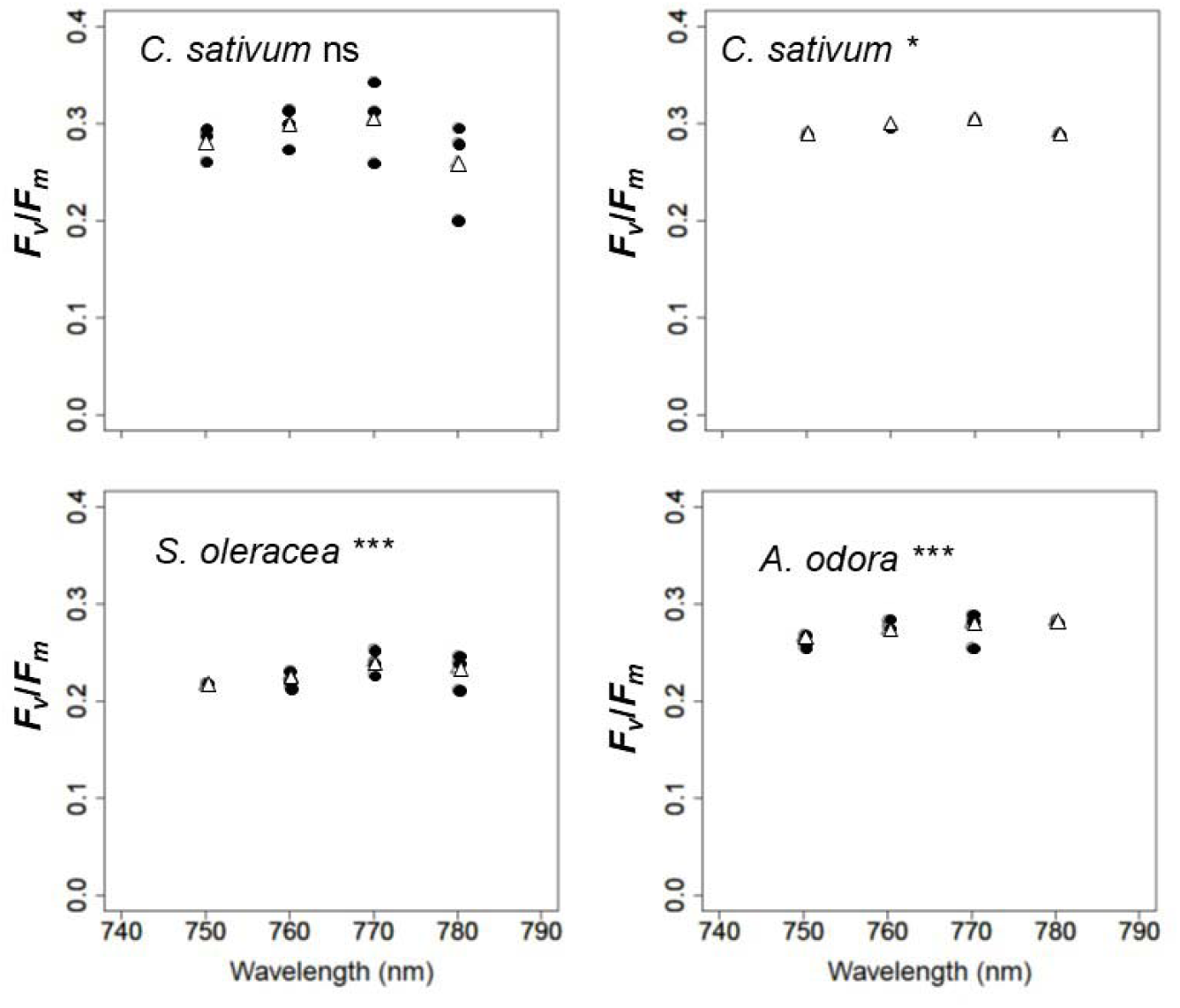
The data used for the fitting and the modelled values. Open circles were the raw data or mean values of the raw data. Solid triangles were modelled *F_v_*/*F_m_* values. * and *** denote that *F*I*_v_*/*F*I*_m_* determined were statistically significant at *P*< 0.05 and *P*< 0.001.

### Is *F*I excited by *F*II?

Using a simple model (Fig. S5), we calculate light absorbed by a thin layer, emission of PSII fluorescence by this layer, re-absorption of this fluorescence by the rest of the leaf, and emission of PSI fluorescence caused by the absorption of PSII fluorescence, in this order. Then, the ratio of the *F*I excited by *F*II to the *F*I directly driven by the blue light will be calculated.

Transmission of the monochromatic light through a pigment solution in a cuvette is expressed by the Lambert-Beer’s law:

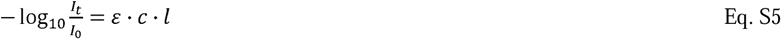

where *I*_0_ and *I_t_* are intensities of the incident light and transmitted light, *l* is the cuvette thickness, *ε* is an absorption coefficient, and *c* is the pigment concentration. Here, we consider the cumulative pigment concentration *C* in mol m^−2^:

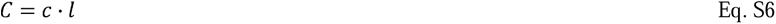

Using a simple model, let us compare PSI fluorescence directly excited by the actinic light and that excited by PS II fluorescence (see Fig. S5).

**Fig. S5.**
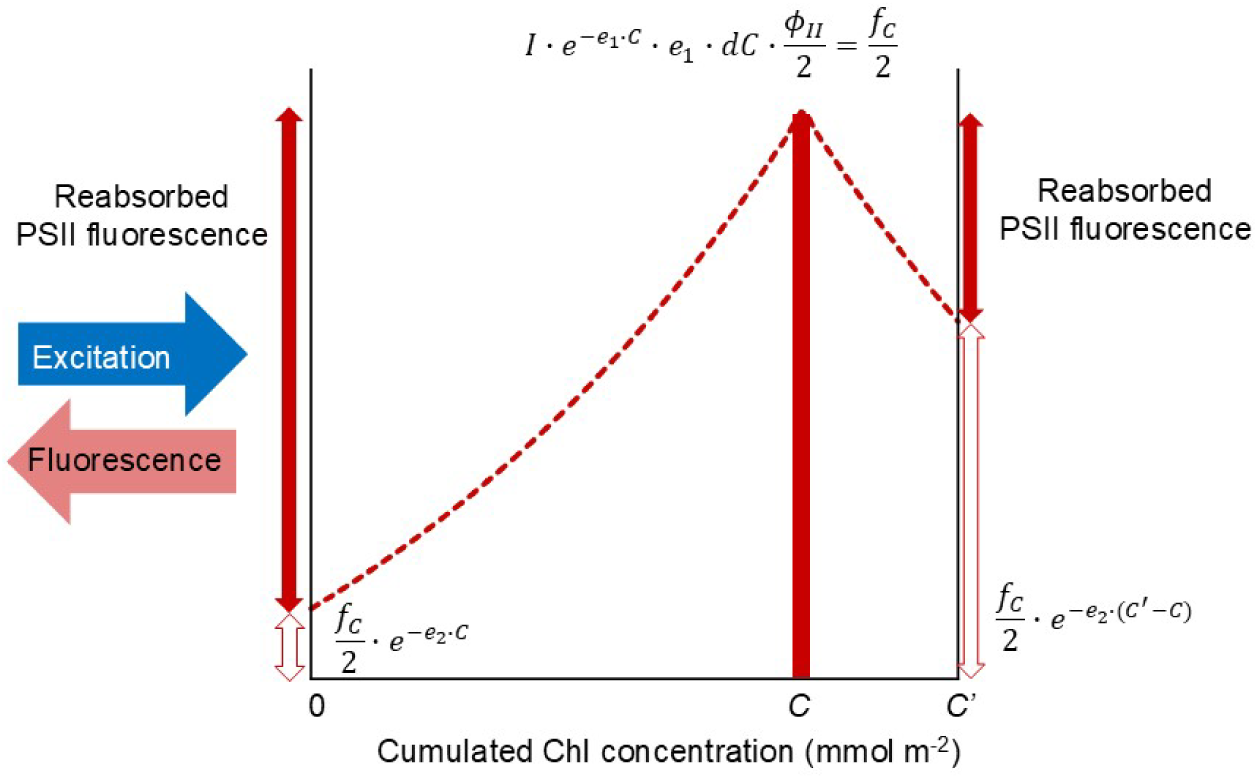
A model leaf used for estimation of PSI fluorescence excited by re-absorbed PSII fluorescence. A thin leaf layer *dC* absorbs light and emits fluorescence. The fluorescence emitted by *dC* is expressed as *f_C_* for simplicity.

Let us consider a thin layer (*dC*) at the cumulated chlorophyll concentration of *C* from the illuminated surface. Light absorption by this thin layer (*dA*) can be written as:

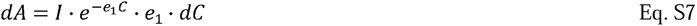

where *e_2_* is equal to 2.30 · ε_1_ and ε_1_ is the absorption coefficient of Chl *a*+*b* at 460 nm (in m^2^ mol^-1^).

Note that ^11^ g, 10 = 2.30. PSII fluorescence emitted by this thin layer can be expressed as:

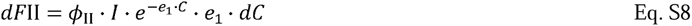

where *Φ_II_* denote PSII fluorescence yield.

Since PSII fluorescence emitted by a thin layer *dC* is absorbed by both sides of the layer, PSII fluorescence re-absorbed in a leaf or solution (0 < X< *C′)* can be expressed as:

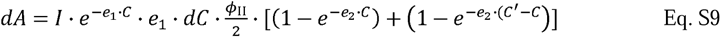

where *e_2_ (=* 2.30 · ε_2_) and ε_2_ *is* the absorption coefficient of Chl *a*+*b* at PSII fluorescence peak (~690 nm). PSI fluorescence excited by PSII fluorescence would not be re-absorbed. Then, PSI fluorescence from *dC* excited by PSII fluorescence and detected at the surface would be half the PSI fluorescence:

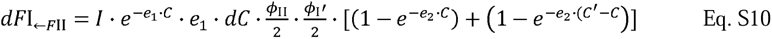

where *Φ_I_*’ denotes the yield of PSI fluorescence excited by PSII fluorescence. On the other hand, PSI fluorescence directly excited by the actinic light, emitted by the thin layer at *C* (*dC*), and detected by the sensor is expressed as:

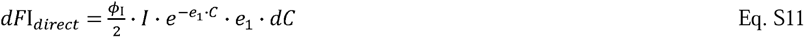

where *Φ_I_* denotes the yield of PSI fluorescence excited by the blue actinic light. Integration from 0 to *C* of these equations give:

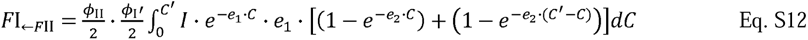

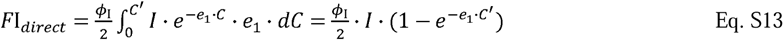

The ratio of these integrals is expressed as:

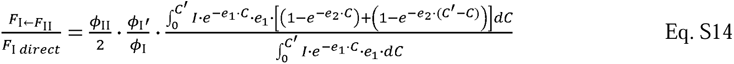

Because we may neglect the effects of PSII fluorescence emitted to the other side, PSII fluorescence emitted from *dC* and reaches the surface can be expressed as:

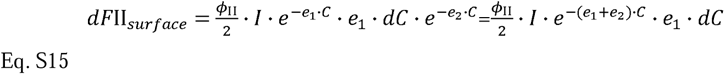

The absorption coefficients *in situ*, *e*_1_ and *e*_2_, may be estimated by examining the effects of the cumulated Chl concentration of the thylakoid suspension on PSI and PSII fluorescence.

PSII fluorescence reaching the surface can be estimated by integrating Eq. S15:

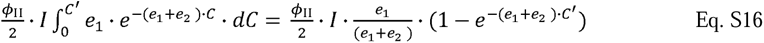

By curve fitting of the dependence of PSI and PSII fluorescence (Eqs. S13 and S16) on *C′*, *e*_ļ_ and *e2* may be estimated (see Fig. S6 for the estimation of *e*_1_).

The ratio (Eq. S14) is a function of PSII and PSI fluorescence yields and absorption coefficients. The ratio increases with *C* (Fig. S9). Since PSII fluorescence little excites PSII fluorescence while preferentially excites PSI, *Φ_I_′* would be twice of *Φ_I_* Then, Eq. A10 may become:

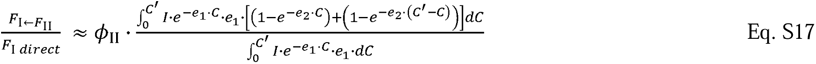

Hence, *Φ_II_* is the key factor. The ratio of the integrals in the realistic *C′* range is at most 1~2 (Fig. S9). Because PSII fluorescence yield excited by blue light, *Φ_II_*, is small (Latimer et al. 1956, Lamb et al. 2018 and references therein), the effects of PSII fluorescence re-absorption on PSI fluorescence should be also small.

**Fig. S6.**
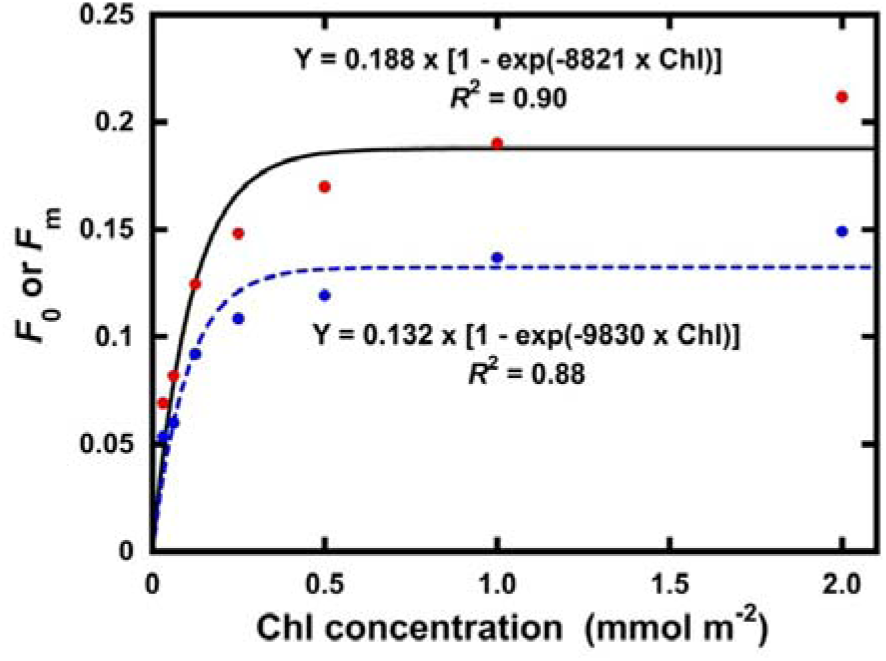
Determination of e1. In this calculation, we assume that no PSI fluorescence is excited by PSII fluorescence.

**Fig. S7.**
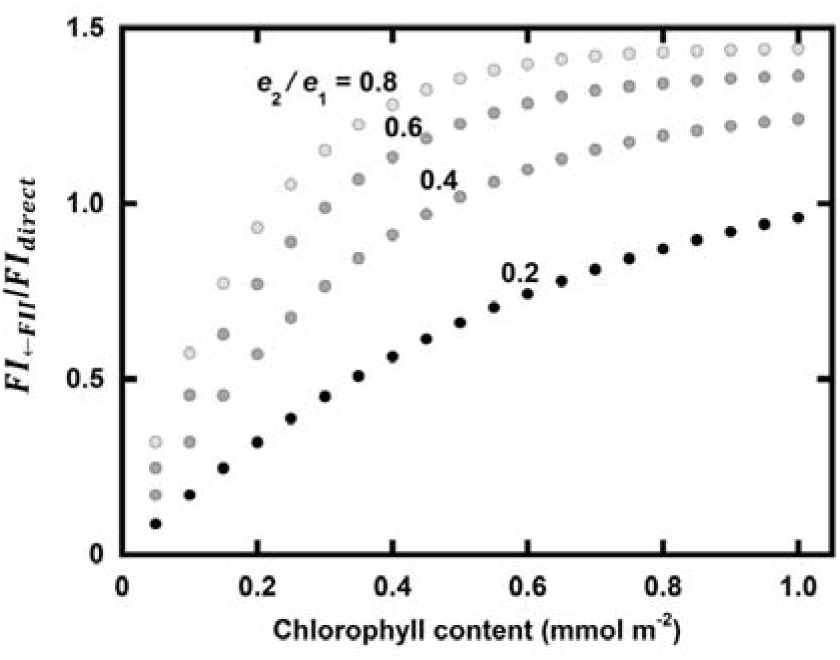
The ratio of the integrations in Eq. S17:

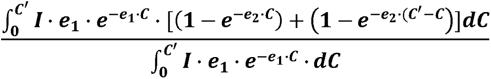

The ratio increases with the increase in *_CF_* For *e _1_,* we used 9000 (namely *ε_1_* of 9000/2.3 = 3913 m^2^ mol^−1^, see Fig. S6). We did not estimate *e*_2_ experimentally. Several possible *e*_2_ values were used (see Fig. S8).

**Fig. S8.**
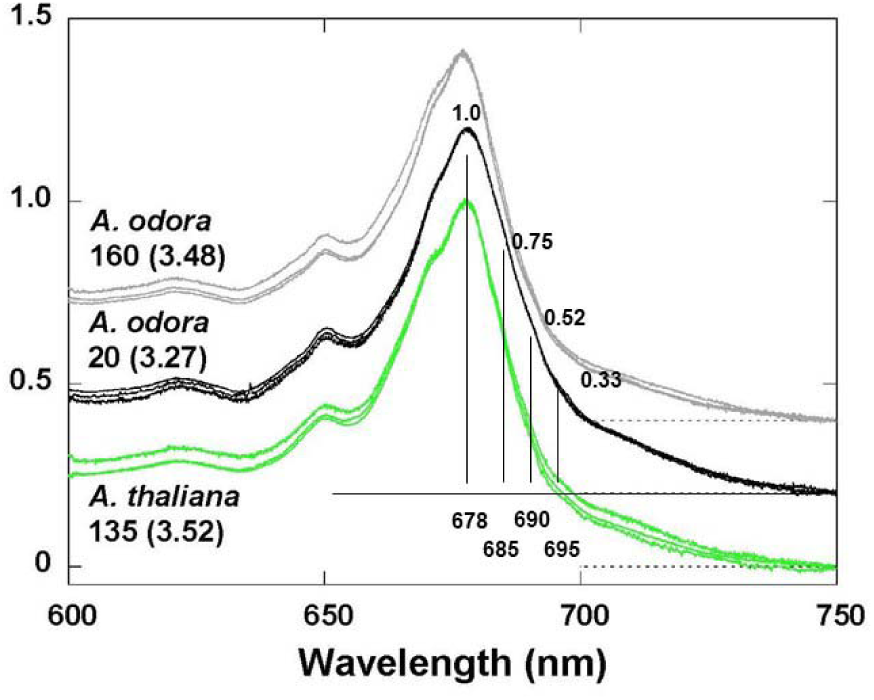
77K absorbance spectra of thylakoid suspensions prepared from the leaves of *A. odora* grown at 160 and 20, and *Arabidopsis thaliana* grown at 135 µmol m^−2^ s^−1^. Assuming the red peak (678 nm) height is similar to that at 460 nm, *e*_2_ at 685, 690 and 695 nm were estimated for low light *A. odora*. The original figure in Terashima et al. (2021) was modified.

**Fig. S9.**
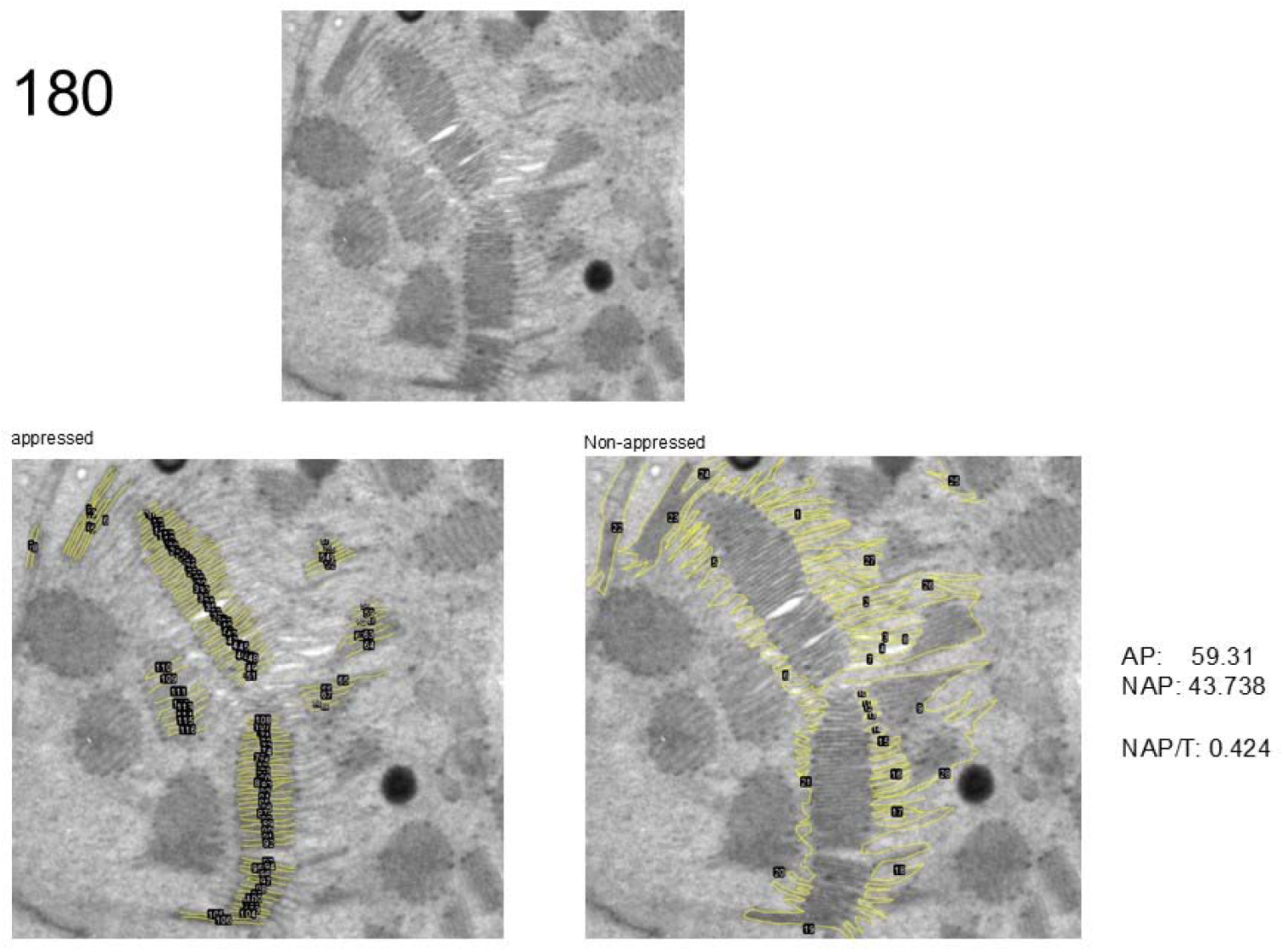
A part of a HL-*A. odora* chloroplast. Determination of the length of non-appressed thylakoid membranes (NAP) and appressed thylakoid membranes (AP). The ratio of NAP to the total length (NAP+AP) was 0.424.

**Fig. S10.**
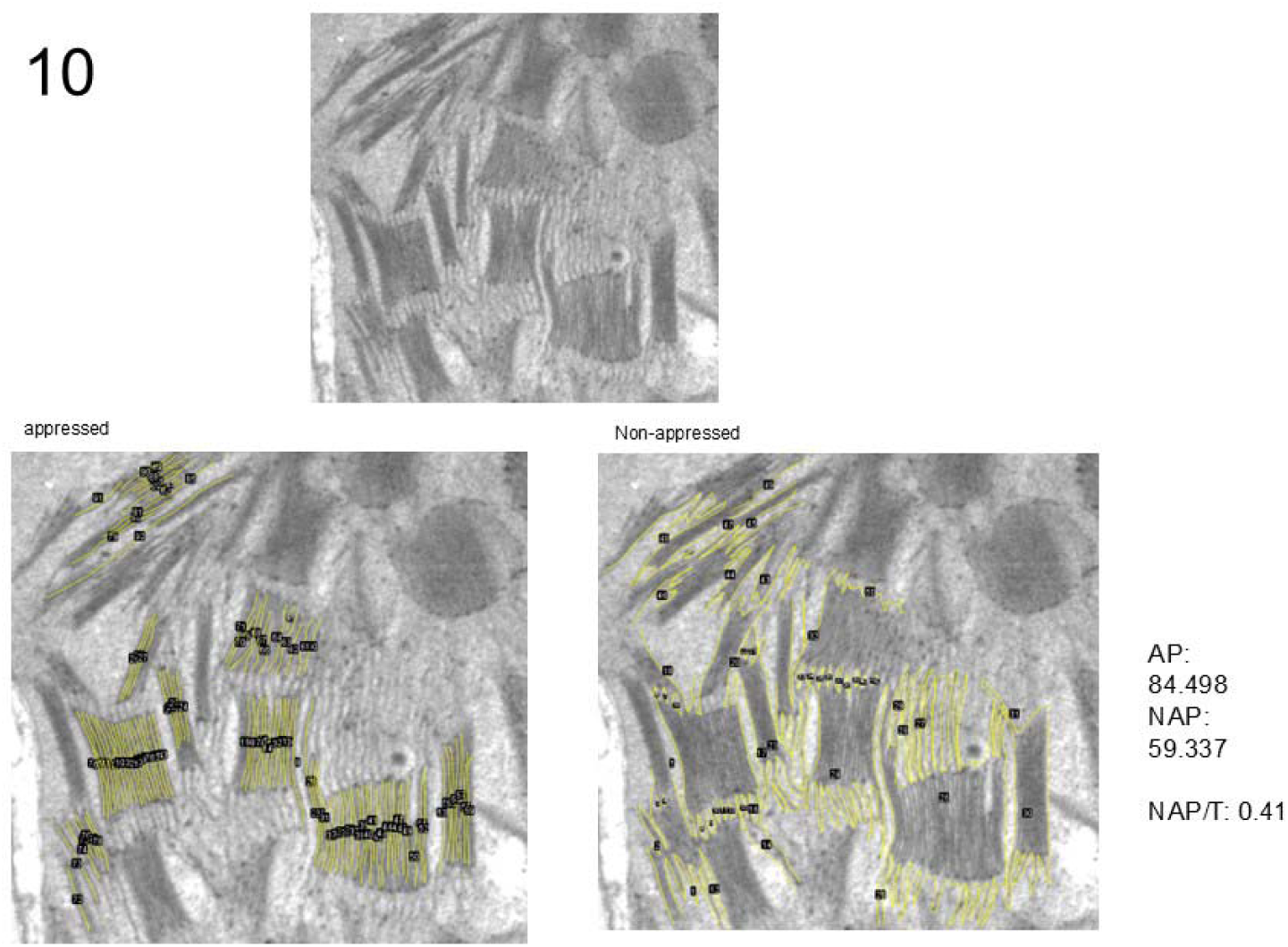
A part of a HL-*A odora* chloroplast. Determination of the length of non-appressed thylakoid membranes (NAP) and appressed thylakoid membranes (AP). The ratio of NAP to the total length (NAP+AP) was 0.41.

